# The lipid droplet protein Jabba promotes actin remodeling downstream of prostaglandin signaling during Drosophila oogenesis

**DOI:** 10.1101/2025.05.09.653087

**Authors:** Jonathon M. Thomalla, Michelle S. Giedt, Roger P. White, Israel J. Wipf, Alicia Shipley, Michael A. Welte, Tina L. Tootle

**Affiliations:** Department of Molecular Biology and Genetics, Cornell University, Ithaca, NY; Department of Biology, University of Iowa, Iowa City, IA; Department of Biology, University of Rochester, Rochester, NY

## Abstract

Growing evidence supports that lipid droplets (LDs) are critical for producing high-quality oocytes. However, the functions of LDs during oocyte development remain largely unknown. Using Drosophila oogenesis as a model, we previously discovered the LD-associated Adipose Triglyceride Lipase (ATGL) promotes actin remodeling necessary for oocyte development by providing the substrate for producing lipid signals termed prostaglandins (PGs). Here we find that Jabba, a LD-associated protein best known for its role in anchoring other proteins to LDs, also promotes PG-dependent actin remodeling. Overexpression of Jabba results in thickened cortical actin and excessive actin bundles, whereas loss of Jabba results in cortical actin breakdown and severely defective actin bundle formation. We find that Jabba regulates actin remodeling independently of ATGL but in conjunction with PG signaling. These data support that there are two PG signaling pathways that promote actin remodeling: one PG pathway that is dependent on ATGL and the other requires Jabba. Overexpression of Jabba rescues the actin defects when PG signaling is lost. Together these data lead to the model that PGs produced independently of ATGL positively regulate Jabba to promote actin remodeling necessary for follicle morphogenesis and the production of a fertilization competent oocyte.

**Significance statement:**

- Across organisms, lipid droplets accumulate during oocyte development and are implicated in fertility. The functions of lipid droplets during oogenesis are poorly understood.
- The authors use the genetic tools and well-characterized process of Drosophila oogenesis to reveal that Jabba, a lipid droplet anchoring protein, is a new downstream effector of prostaglandin signaling and promotes actin remodeling necessary for producing a fertilization competent oocyte.
- The results extend prior studies connecting lipid droplet proteins, prostaglandins, and actin remodeling, providing insight into how these critical conserved factors contribute to high-quality oocytes.

## INTRODUCTION

A key component of fertility and reproduction is the production of an oocyte that is capable of being fertilized. Oocytes are large cells and often contain substantial stores of nutrients to support the proliferative needs of the eventual embryo (Sturmey *et al*., 2009; Ami *et al*., 2011; Dunning *et al*., 2011; Welte, 2015a; Brusentsev *et al*., 2019; Kilwein *et al*., 2023a). Building an oocyte is therefore a metabolically demanding process, and defective metabolism can lead to severe reductions in fertility and oocyte quality, both in model organisms (Buszczak *et al*., 2002; Jungheim *et al*., 2010; Prates *et al*., 2014) and in humans (Robker, 2008; Cardozo *et al*., 2011). For example, obesity is one underlying contributor to poor oocyte quality and suboptimal in vitro fertilization outcomes in humans (Wattanakumtornkul *et al*., 2003; Styne-Gross *et al*., 2005; Robker, 2008). Thus, understanding the metabolic parameters that define oocyte quality and support peak reproductive success may lead to the development of new treatments or diagnostics for female infertilities, especially those arising from errors in oogenesis.

One key regulator of both oocyte quality and metabolism is lipid droplets (LDs), the cellular organelles for fat storage (Teixeira *et al*., 2003; Yang *et al*., 2010; Dunning *et al*., 2014; Brusentsev *et al*., 2019). LDs are unique organelles with a core of neutral lipids, such as triacylglycerides (TAGs) and sterol esters (SEs) (Walther and Farese, 2012; Hashemi and Goodman, 2015); this core is surrounded by a phospholipid monolayer decorated by a variety of proteins, many of which are involved in lipid metabolism (Welte, 2015b; Bersuker *et al*., 2018; Olzmann and Carvalho, 2019). Indeed, in many cell types, LDs are critical regulators of energy homeostasis and lipid metabolism. In addition, LDs have important roles in cell signaling and regulation of specific proteins (Welte, 2015b; Welte and Gould, 2017); proteomic studies on LDs identified proteins required for DNA damage repair, chromatin assembly, cell signaling, and cytoskeletal regulation (Cermelli *et al*., 2006; Bersuker *et al*., 2018).

LDs are present during oogenesis in many species, accumulating to varying degrees in organisms ranging from flies to mammals (Buszczak *et al*., 2002; Teixeira *et al*., 2003; Ambruosi *et al*., 2009; Dunning *et al*., 2014; Aizawa *et al*., 2024). These LDs are likely a source of energy for oogenesis and the future embryo (Dunning *et al*., 2010; Yang *et al*., 2010; Dunning *et al*., 2014; Brusentsev *et al*., 2019); however, the non-metabolic roles of LDs for oocyte development are generally uncharacterized, and the mechanisms by which LDs and their associated proteins contribute to oocyte development remain largely unknown.

Drosophila oogenesis is an ideal model system for addressing how LD regulation during oogenesis contributes to fertility. Drosophila eggs are laid and develop externally, and thus the accumulation and regulation of LDs during oogenesis is critical for the future embryo (Welte, 2015a; Kilwein *et al*., 2023a). Within the abdomen of the female fly are two ovaries, each of which is composed of 15-20 ovarioles. Ovarioles are chains of follicles, also termed egg chambers, that are arranged in order of maturity; there are fourteen stages (S) of follicle development. Each follicle is composed of a layer of somatic cells termed follicle cells which surround 15 germline-derived nurse cells and the oocyte (Giedt and Tootle, 2023). LDs are present in small numbers in early stages of oogenesis but begin to accumulate massively within the nurse cells during S9; by S10B, LDs are densely packed throughout the nurse cell cytoplasm (Buszczak *et al*., 2002; Teixeira *et al*., 2003; Stephenson *et al*., 2021). During the next stage of oogenesis, the nurse cells contract and transfer LDs, as well as the rest of their cytoplasm, to the oocyte through intracellular bridges, in a process termed nurse cell dumping (Wheatley *et al*., 1995; Guild *et al*., 1997; Huelsmann *et al*., 2013). Follicle LDs store both TAGs and SEs (Heier *et al*., 2021; Giedt *et al*., 2023), and conserved TAG biosynthesis enzymes, such as DGAT1/Midway, are essential for LD accumulation and oogenesis (Buszczak *et al*., 2002; Parra-Peralbo and Culi, 2011). Further, LD-associated proteins in nurse cells have recently been ascribed developmental roles; namely, they control the oocyte’s histone levels and regulate nurse cell actin structures to facilitate nurse cell dumping (Teixeira *et al*., 2003; Cermelli *et al*., 2006; Stephenson *et al*., 2021; Giedt *et al*., 2023).

LDs are necessary for the oocyte to provide high levels of histones to the early embryo (Cermelli *et al*., 2006; Li *et al*., 2012; Stephenson *et al*., 2021). Drosophila oocytes have large cytoplasmic stores of histones that promote chromatin assembly in the early embryo (Li *et al*., 2012; Tirgar *et al*., 2023). For the histones H2A, H2B, and H2Av, these stores are present on LDs, attached via the anchoring protein Jabba (Li *et al*., 2012). Jabba dependent LD-sequestration of histones is already detected in S9 nurse cells and may promote the transfer of these histones to the oocyte. In the oocyte itself, Jabba protects the histones from degradation (Stephenson *et al*., 2021).

In S10B nurse cells, LDs also promote the production of prostaglandins (PGs) which in turn control the remodeling of the actin cytoskeleton (Giedt *et al*., 2023). The TAGs in follicle LDs contain many different types of fatty acids, including the poly-unsaturated arachidonic acid (AA). AA can be released from LDs via Adipose Triglyceride Lipase (ATGL/Brummer) and is converted, via the cyclooxygenase-like enzyme dCOX1 (also known as Pxt) and downstream synthases, to PGF_2α_, a small lipid signaling molecule from the PG family. PGF_2α_ in turn signals to promote actin remodeling (Tootle and Spradling, 2008; Groen *et al*., 2012; Spracklen *et al*., 2014; Spracklen *et al*., 2019; Giedt *et al*., 2023): The cortical actin at the periphery of each nurse cell thickens and contracts, and actin bundles form at the periphery and extend inward toward the nuclei. Proper remodeling of both types of actin is critical for nurse cell dumping (Wheatley *et al*., 1995; Guild *et al*., 1997; Huelsmann *et al*., 2013) and thus essential for both follicle morphogenesis and the production of a fertilization competent, high-quality oocyte.

Jabba was previously shown to affect histone levels in the cytoplasm and nuclei and to prevent inappropriate interactions between LDs and glycogen granules (Johnson *et al*., 2018; Stephenson *et al*., 2021; Kilwein *et al*., 2023a; Kilwein *et al*., 2023b). In this study, we uncover a new function for Jabba: loss or overabundance of Jabba alters actin remodeling in S10B nurse cells. Using genetic interaction studies and rescue experiments, we find that Jabba regulates the actin cytoskeleton in conjunction with PG signaling, but in an unexpected manner: it acts independently of ATGL and downstream of PG production. Our data supports that there are at least two LD-PG signaling pathways that promote actin remodeling necessary for follicle development; one PG pathway is dependent on ATGL and the other involves Jabba. Together these data lead to the model that PGs produced independently of ATGL positively regulate Jabba to promote actin remodeling necessary for follicle morphogenesis and the production of a healthy mature oocyte.

## RESULTS

### Overexpression of the LD protein Jabba results in excessive actin assembly

During Drosophila oogenesis, the LD-associated protein ATGL is a critical regulator of actin cytoskeletal remodeling (Giedt *et al*., 2023). We therefore asked if the proper regulation of actin remodeling during follicle development also involves other LD-associated proteins. A promising candidate is Jabba as it was previously reported that Jabba knockdown in the female germline disrupts lipid metabolism (McMillan *et al*., 2018). In previous studies, we generated genomic transgenes that make it possible to proportionally increase Jabba protein levels in follicles (Johnson *et al*., 2018; Stephenson *et al*., 2021). We therefore compared S10B follicles with three different *Jabba* gene dosages: wild type (*1x Jabba*) or otherwise wild-type lines with one (*1.5x Jabba*) or two (*2x Jabba*) copies of the transgene.

Fixed follicles were stained with phalloidin to detect filamentous actin (F-actin) and analyzed by confocal microscopy. We analyzed two populations of actin in S10B nurse cells: a meshwork of cortical actin directly beneath the plasma membrane, around the entire cell periphery; and actin bundles that originate at the periphery and project toward the nuclei (Figure 1A-A’). We found that both populations of F-actin are increased in a *Jabba* dosage-dependent manner: in follicles with higher Jabba levels, the cortical actin appeared thicker, and actin bundles tended to be more abundant (Figure 1B-C’). To quantify these changes in the actin cytoskeleton, confocal image stacks of S10B follicles were scored for two criteria: do they display excessively thickened cortical actin and/or increased numbers of actin bundles? While only a small fraction of wild-type follicles (4%) showed excessive actin accumulation, 38% of *1.5x Jabba* follicles and 67% of *2x Jabba* follicles exhibit overabundant actin (Figure 1D). These data suggest that Jabba can promote actin assembly, at least when overexpressed.

**Figure 1.**
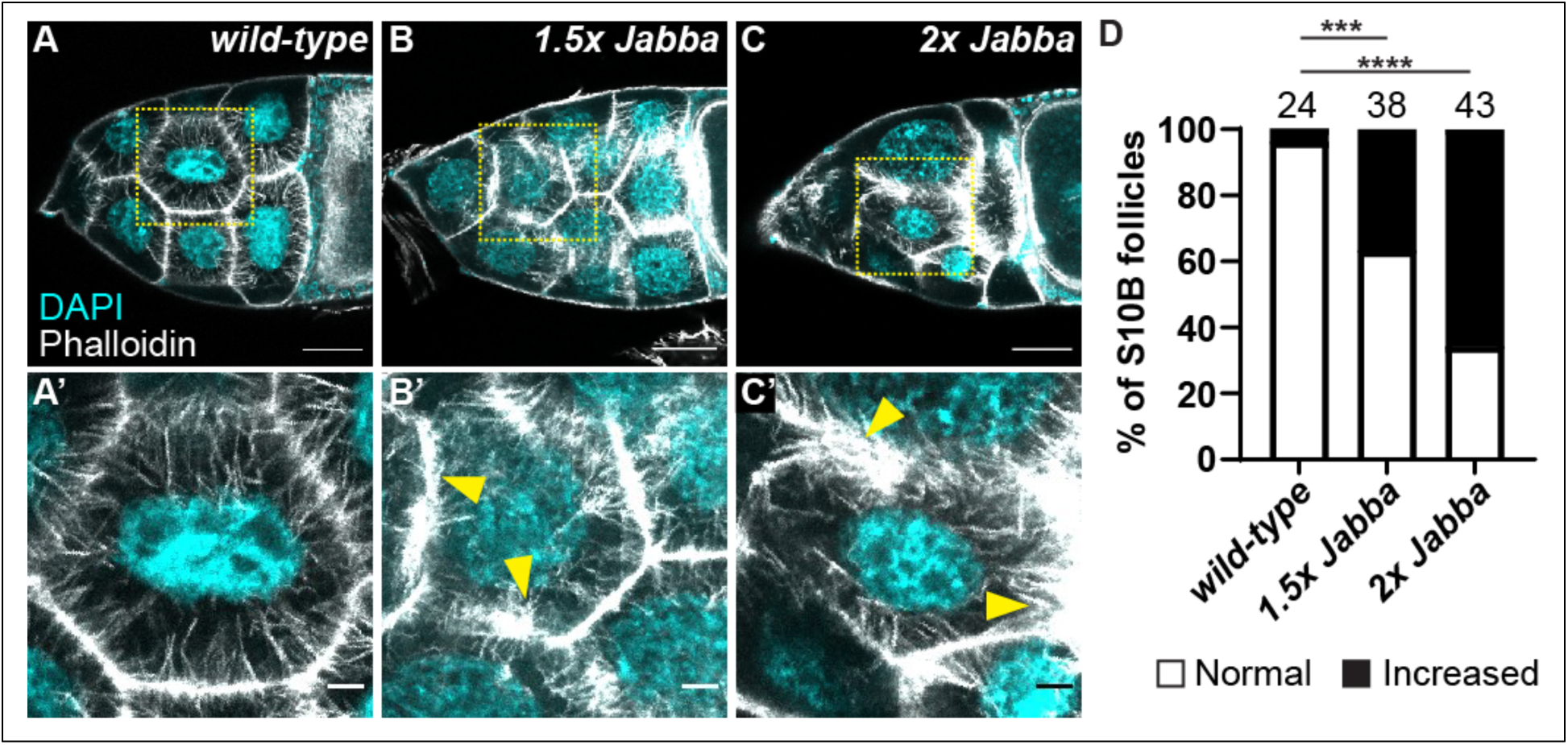
Jabba promotes actin assembly. (A-C’) Maximum projection of three confocal slices of S10B follicles stained for F-actin (phalloidin) in white, and DNA (DAPI) in cyan. Yellow dashed boxes in A-C indicate zoomed in regions in A’-C’. Yellow arrowheads indicate regions of increased actin. A black box was added behind the label C’ to improve visualization. Scale bars in A-C = 50µm and A’-C’ = 10µm. \*\*\**p<0.001, ****p<0.0001,* Pearson’s chi-squared test. (A, A’) *wild type* (*yw*). (B, B’) *1.5x Jabba* (one copy of the genomic *Jabba* transgene in a wild-type background). (C, C’) *2x Jabba* (two copies of the genomic *Jabba* transgene in a wild-type background). (D) Graph quantifying the prevalence of normal versus thickened cortical and/or bundled actin. Wild-type follicles have a cortical actin boundary and evenly spaced actin bundles around the nurse cell periphery (A-A”, D). Additional copies of Jabba (*1.5x or 2x Jabba*) result in a dosage-dependent appearance of thicker cortical actin and either thicker or more numerous actin bundles (B-D). Follicles exhibiting the strongest increases in F-actin are sometimes accompanied by cortical actin breakdown.

### Loss of Jabba impairs actin remodeling

To determine whether Jabba promotes actin assembly under normal conditions, we examined the actin cytoskeleton in flies homozygous for either of two strong loss-of-function *Jabba* alleles: *Jabba^DL^* deletes most of the Jabba coding region; *Jabba^z101^* lacks the main promoter of the *Jabba* locus (Li *et al*., 2012). No Jabba protein is detectable in the embryos or oocytes of either genotype (Li *et al*., 2012). In contrast to the uniform cortical actin and straight actin bundles in wild-type S10B nurse cells (Figure 2A-A’), both mutants exhibited various actin defects, including decreased or absent bundles and localized breakdown of the cortical actin (Fig. 2B-B’). To quantify the frequency and severity of actin defects, we used a previously developed method (Giedt *et al*., 2023), where we analyzed confocal stacks of S10B follicles for sparse, missing, or collapsed actin bundles and disrupted cortical actin. Actin bundle defects and cortical actin defects were scored separately by binning them into four categories giving a score of 0-3, with 0 indicating no defects (aka normal actin remodeling). The scores for actin bundle and cortical actin defects were then added together to give an Actin Defect Index (ADI) of 0-6. The ADI scores were binned into three categories: normal (scored 0-1), mild (scored 2-3), and severe (scored 4-6). The frequency of actin bundle defects increased from ∼30% in the wild type follicles to ∼70% upon loss of Jabba (Figure 2D). Similarly, none of the wild-type S10B follicles displayed disruptions in their cortical actin, but 60% of *Jabba* mutant follicles did (Figure 2F). Overall, total actin remodeling was largely normal in wild-type follicles (96% normal ADI) but drastically altered in *Jabb*a mutants (only 40% normal ADI; Figure 2G). Thus, actin remodeling is severely defective in the absence of Jabba. Similar but milder actin defects are observed when Jabba is knocked down in the germline using RNAi (Supplemental Figure S1), suggesting Jabba likely acts within the nurse cells to regulate actin remodeling. Together with our overexpression studies, we conclude that actin assembly is sensitive to Jabba levels and that Jabba promotes actin assembly.

**Figure 2.**
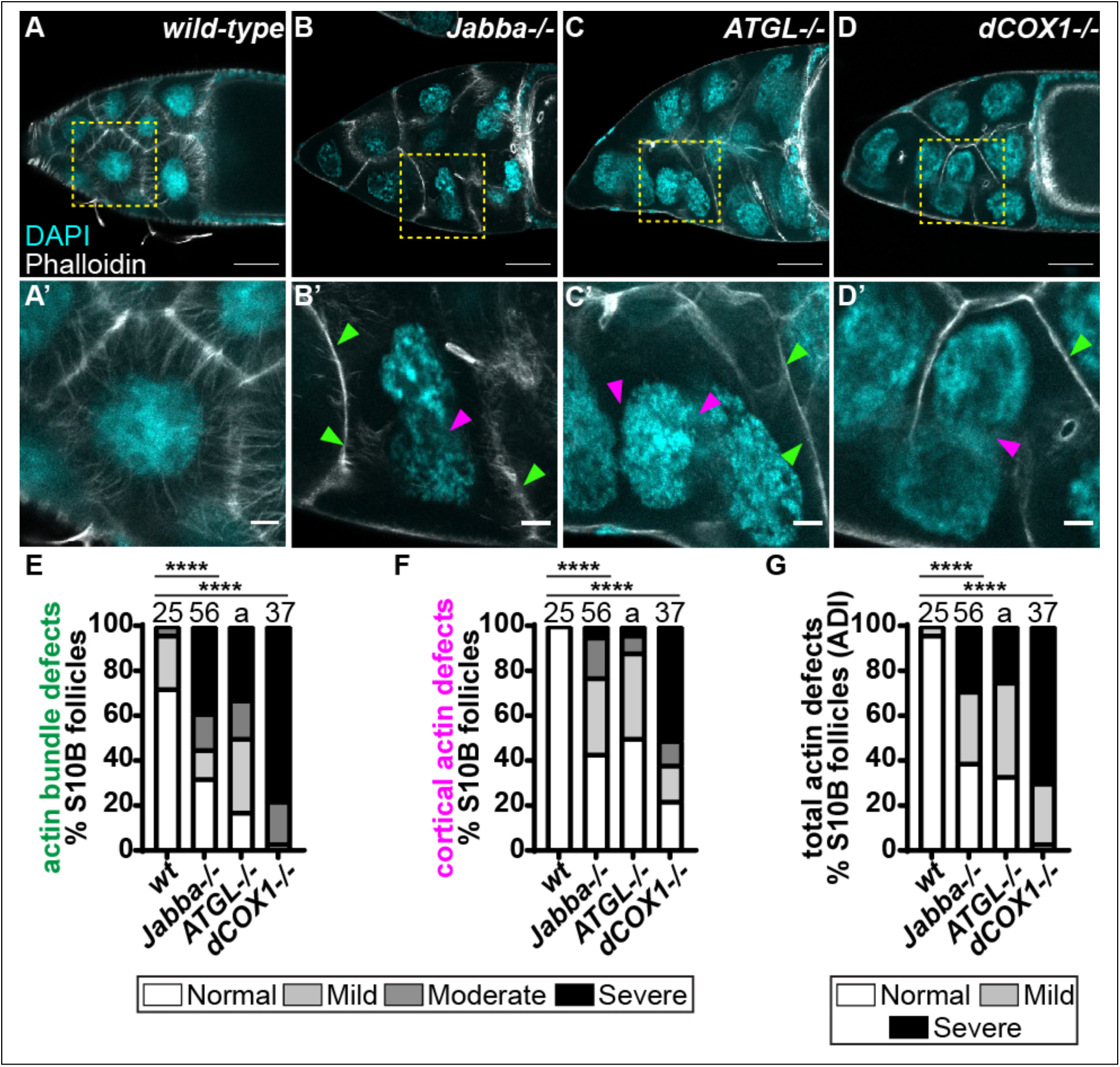
Jabba is required for normal actin remodeling. (A-D’) Maximum projection of three confocal slices of S10B follicles stained for F-Actin (phalloidin) in white, and DNA (DAPI) in cyan. Yellow dashed boxes in A-D indicate zoomed in regions in A’-D’. Arrowheads indicate examples of defective actin bundling (green) and disrupted cortical actin (magenta). Scale bars in A-D = 50µm and A’-D’ = 10µm. (A, A’) *wild type* (*yw*). (B, B’) *Jabba-/-* (*Jabba^z101^/Jabba^z101^*). (C, C’) *ATGL-/-* (*bmm^1^/bmm^1^*). (D, D’) *dCOX1-/-* (*pxt^f01000^/ pxt^EY03052^*). (E-G) Graphs quantify the actin phenotypes of the following genotypes: *wild-type* = *yw. Jabba-/-* = *Jabba^zl01^/Jabba^zl01^* and *Jabba^DL^/Jabba^DL^*. *ATGL-/-* = *bmm^1^/bmm^1^*. *dCOX1-/-* = *pxt^f01000^/pxt^f01000^*and *pxt^EY03052^/pxt^EY03052^*. Actin defects were quantified by scoring the penetrance of actin bundle and cortical actin defects into one of four categories: normal or mild, moderate or severe defects. Scores were summed and the total binned into one of three total actin defects (ADI) categories: normal, mild defects, or severe defects. For a detailed description of the quantification refer to Materials and Methods. The “a” indicates previously published data (*p<0.0001,* (Giedt et al., 2023)). *\*\*\*\*p<0.0001,* Pearson’s chi-squared test. In wild-type S10B follicles, actin bundles extend from the nurse cell periphery to the nucleus, and the cortical actin is thickened relative to earlier stages (A, A’). Actin bundles fail to form or form improperly, and cortical actin is disrupted upon loss of Jabba (B, B’), ATGL (C, C’) or dCOX1 (D, D’). Loss of Jabba, ATGL, or dCOX1 significantly increases the frequency of actin bundle (E) and cortical actin defects (F) compared to wild-type S10B follicles. Compared to wild type, mutants in *Jabba*, *ATGL*, or *dCOX1* display a significantly lower percentage of normal follicles and a higher percentage of follicles exhibiting severe total actin defects (G).

### Jabba and ATGL work in separate pathways to regulate actin remodeling

The S10B actin defects in *Jabba* mutants resemble those we previously described upon loss of either ATGL, a triglyceride lipase, or dCOX1, the key enzyme in PG synthesis (Tootle and Spradling, 2008; Spracklen *et al*., 2014; Giedt *et al*., 2023). Both *ATGL* and *dCOX1* mutants exhibit defects in or loss of actin bundles as well as breakdown of cortical actin (Figure 2C-G). This combination of phenotypes is unusual as most known actin regulators affect either cortical actin or actin bundles, but not both (Wheatley *et al*., 1995; Buszczak and Cooley, 2000). Similar phenotypes suggest Jabba, ATGL, and dCOX1 may all function through a common pathway to regulate actin remodeling. Indeed, we previously found that ATGL regulates the release of LD-housed AA, the substrate for PG production; PG signaling then drives actin remodeling (Giedt *et al*., 2023). We therefore tested whether Jabba affects actin remodeling via the ATGL-AA pathway.

We first performed a dominant genetic interaction assay between Jabba and ATGL using the actin defect quantification method described above (Giedt *et al*., 2023). This assay relies on heterozygosity for mutations in *Jabba* or *ATGL* having only very modest effects on actin remodeling, resulting in largely normal actin structures. Both genotypes had ∼40% of follicles with actin bundle defects, with severe defects being rare (7% in *ATGL*-/+ follicles; none detected in *Jabba*-/+ follicles; Figure 3 A-B, D). Disrupted cortical actin was observed at a frequency of 16% (0% severe) in *Jabba*-/+ and 12% (7% severe) in *ATGL-/+* follicles (Figure 3E). Examining double heterozygotes for *Jabba* and *ATGL* can then be used to interrogate whether Jabba and ATGL function in the same pathway. If they are, then double heterozygotes will exhibit a synergistic increase in severe actin defects. Conversely, if they function in separate pathways, then the actin defects in the double heterozygotes will remain low or be additive of what is observed in the two single heterozygotes. Using this assay, we previously found that actin defects in double heterozygotes of ATGL and dCOX1 are synergistic (Giedt *et al*., 2023) and thus that ATGL and dCOX1 act in the same pathway. The double heterozygotes of Jabba and ATGL showed a higher frequency of severe actin bundle defects (20% severe), but the total frequency of defective bundles was 45%, comparable to that seen in the single heterozygotes (Figure 3D). Likewise, defective cortical actin was seen in 18% (2% severe) of double heterozygotes (Figure 3E), very similar to the single heterozygotes. Overall, total actin remodeling was very similar for *Jabba*-/+ (24% defective ADI), *ATGL-/+* (19% defective ADI), and *Jabba*-/+; *ATGL*-/+ (32% defective ADI), which points to an additive effect of these genes rather than a synergistic interaction (Figure 3F).

**Figure 3.**
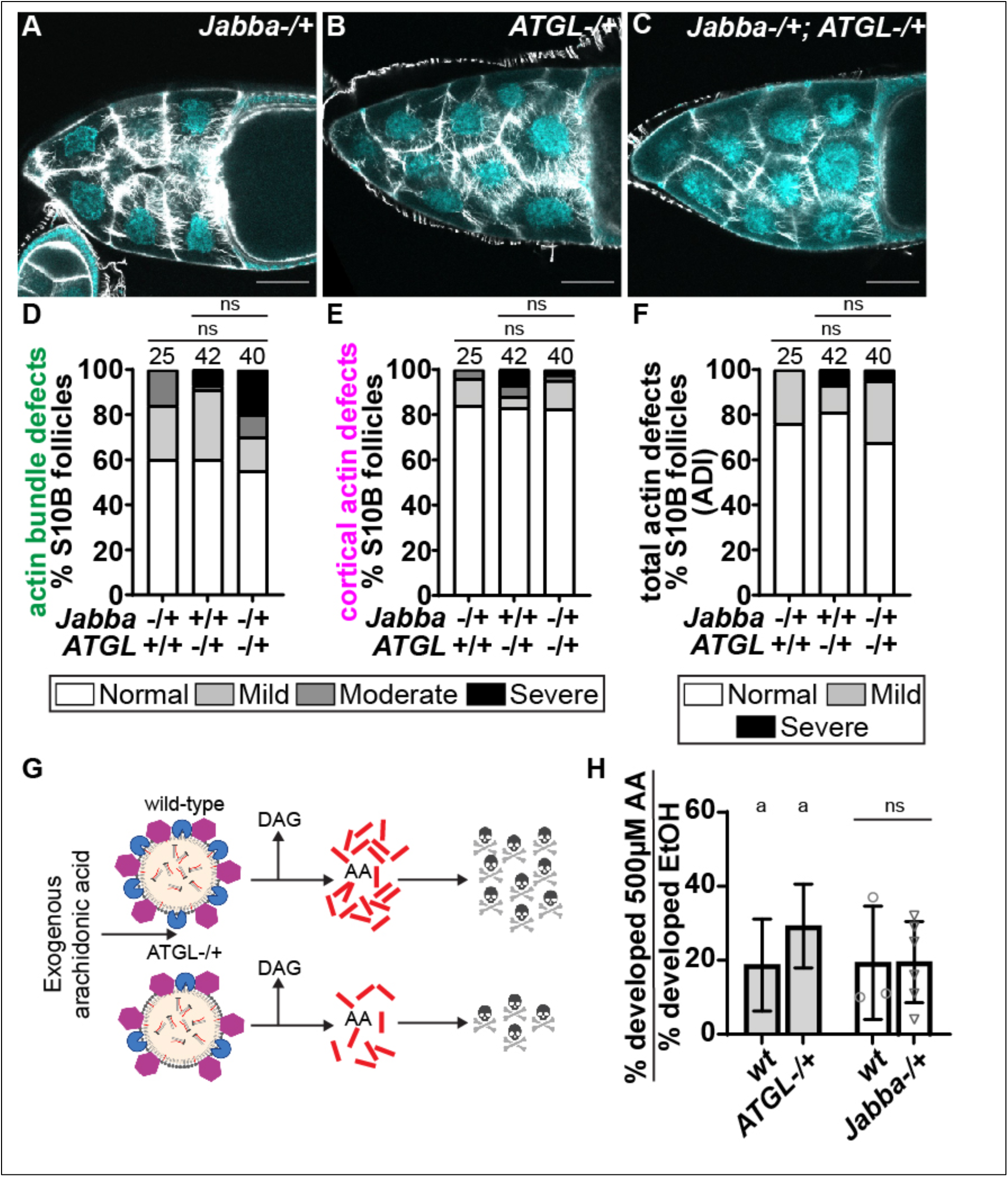
Jabba regulates actin remodeling independently of ATGL. (A-C) Maximum projections of three confocal slices of S10B follicles stained for F-Actin (phalloidin) in white, and DNA (DAPI) in cyan. Scale bars = 50μm. (A) *Jabba-/+* (*Jabba^zl01^/+*). (B) *ATGL-/+* (*bmm^1^/+*). (C) *Jabba-/+; ATGL-/+* (*Jabba^z101^/+; bmm^1^/+*). (D-F) Graphs quantify the actin phenotypes of the following genotypes: *Jabba-*/+ = *Jabba^zl01^/+*; *ATGL-/+ = bmm^1^/+*; *Jabba-/+; ATGL-/+ = Jabba^zl01^/+; bmm^1^/+*. Actin defects were quantified by scoring the penetrance of actin bundle and cortical actin defects into one of four categories: normal or mild, moderate, or severe defects. Scores were summed and the total binned into one of three total actin defects (ADI) categories: normal, mild defects, or severe defects. For a detailed description of the quantification refer to Materials and Methods. ns *p>0.05,* Pearson’s chi-squared test. (G) Schematic of the rationale behind arachidonic acid (AA, red lines) treatment of S10B follicles (made in BioRender). On the LDs, ATGL is shown in blue and Jabba is magenta. DAG is diacylglycerol and the skull and crossbones indicate follicle death. (H) Graph of the ratio of the percentage of S10B follicles developing in the AA medium to the percentage developing in the control medium for the given genotypes; WT (*yw*), *ATGL* (*bmm^1^/+*), *Jabba* (*Jabba^zl01^/+* and *Jabba^DL^/+).* The letter “a” indicates previously published data (*p<0.05,* (Giedt et al., 2023)). Error bars = SD. ns *p>0.05,* two-tailed, paired t-test. Heterozygosity for mutations in *Jabba* (*Jabba-/+*) or *ATGL* (*ATGL*-/+) or reduction of both via double heterozygotes (*Jabba*-/+; *ATGL*-/+) result in largely normal cortical and bundled actin (A-F). Excess exogenous AA is incorporated into LDs which can be liberated by ATGL; high levels of AA is toxic to follicles and prevents follicle maturation in the IVEM assay (G). We previously showed that the toxicity of high levels of AA on follicle development is suppressed by reducing ATGL levels by heterozygosity (left side of H, (Giedt et al., 2023)). In contrast, heterozygosity for Jabba does not relieve AA-induced toxicity (right side of H).

Next, we tested if Jabba regulates AA trafficking. In S10B follicles, exogenous AA exhibits a dose-dependent toxicity which is ameliorated by rapid sequestration of AA into LDs (Figure 3G, (Giedt *et al*., 2023)). This toxicity can be used to study how well LDs sequester or release AA. We first confirmed that AA is still incorporated into LDs in the absence of Jabba (Supplemental Figure S2). We then assessed the effect of exogenous AA on follicle development, using an *in vitro* follicle maturation (IVEM) assay, in which S10B follicles can mature to S14 in culture (Tootle and Spradling, 2008; Spracklen and Tootle, 2013). Specifically, exposure to a high dose of AA (500µM) severely impairs wild-type S10B follicle development in culture (Giedt *et al*., 2023). This AA-induced toxicity is suppressed by reducing the level of ATGL (Figure 3G-H), presumably because ATGL normally releases AA from LD triglycerides and thus increases the pool of free, toxic AA (Giedt *et al*., 2023). In contrast, when we reduced Jabba levels (*Jabba-/+*), AA toxicity was unchanged (Figure 3H), suggesting that – unlike ATGL – Jabba does not mediate AA release from LDs. Together with our genetic interaction assay, we conclude that Jabba and ATGL promote actin remodeling via distinct pathways.

### Jabba functions in a pathway with dCOX1

Even though loss of Jabba and ATGL leads to similar actin defects, we find no evidence that the two LD proteins act in the same pathway. As ATGL regulates actin via PG signaling, our results suggest Jabba controls actin via a novel, PG-independent pathway. To directly test this idea, we performed our dominant genetic interaction assay between Jabba and dCOX1. Follicles heterozygous for mutations in *Jabba* or *dCOX1* exhibited actin bundle defects at a frequency of 27% (0% severe) and 37% (7% severe), respectively (Figure 4A-B, D). Disrupted cortical actin occurred in 10% of *Jabba*-/+ (0% severe) and *dCOX1*-/+ (3% severe) follicles (Figure 4A-B, E). Surprisingly and contrary to our hypothesis, the double heterozygotes (*Jabba*-/+; *dCOX1*-/+) displayed severe actin defects. 95% of follicles had missing or stunted actin bundles (∼50% severe) and ∼70% had disrupted cortical actin (∼20% severe; Figure 4C-E). Overall, total actin remodeling was largely normal for Jabba heterozygotes (90% normal ADI) and dCOX1 heterozygotes (80% normal ADI, Figure 4F). In contrast, only 11% of the follicles from the double heterozygotes (*Jabba-/+; dCOX1-/+*) had a normal ADI and the frequency of the severe class increased almost tenfold (Figure 4F). This synergistic increase in actin defects in the double heterozygotes suggests Jabba and dCOX1 act in the same pathway to regulate actin remodeling. Based on the lack of interaction between Jabba and ATGL (Figure 3), this finding suggests that PGs regulate actin remodeling via two distinct pathways: one dependent on ATGL-AA and one dependent on Jabba.

**Figure 4.**
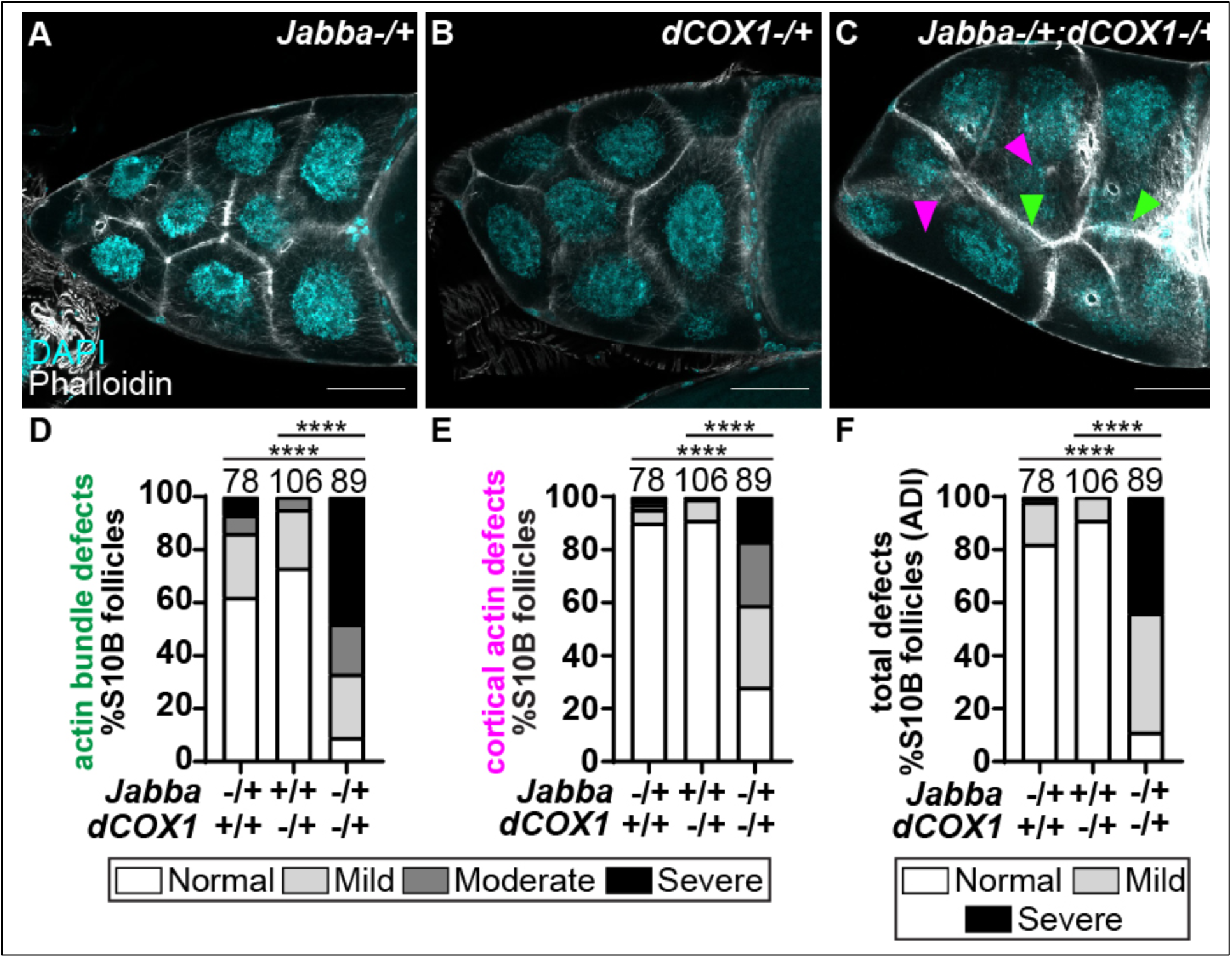
Jabba acts in the same pathway as PGs to promote actin remodeling. (A-C) Maximum projections of three confocal slices of S10B follicles stained for F-Actin (phalloidin) in white, and DNA (DAPI) in cyan. Arrowheads indicate examples of defective actin bundling (green) and disrupted cortical actin (magenta). Images were brightened by 30% to increase clarity. Scale bars = 50μm. (A) *Jabba-/+* (*Jabba^z101^/+*). (B) *dCOX1-/+* (*pxt^EY03052^/+*). (C) *Jabba-/+; dCOX1-/+* (*Jabba^z101^/+; pxt^f01000^/+*). (D-F) Graphs quantify the actin phenotypes of the following genotypes: *dCOX1-/+* = *pxt^f01000^/+* and *pxt^EY03052^/+; Jabba-/+ = Jabba^z101^/+* and *Jabba^DL^/+*; *Jabba-/+; pxt-/+* = *Jabba^z101^/+; pxt^f01000^/+, Jabba^z101^/+; pxt^EY03052^/+, Jabba^DL^/+; pxt^f01000^/+,* and *Jabba^DL^/+; pxt^EY03052^/+.* Actin defects were quantified by scoring the penetrance of actin bundle and cortical actin defects into one of four categories: normal or mild, moderate, or severe defects. Scores were summed and the total binned into one of three total actin defects (ADI) categories: normal, mild defects, or severe defects. For a detailed description of the quantification refer to Materials and Methods. *\*\*\*\*p<0.0001,* Pearson’s chi-squared test. Cortical actin is intact, and actin bundles are straight and arranged around the nurse cell periphery in *Jabba-/+* (A) and *dCOX1-/+* (B) S10B follicles. In contrast, in *Jabba-/+; dCOX1-/+* follicles, actin bundles are absent, sparse, or stunted and there are instances of cortical actin breakdown (C). Quantification reveals a significant increase in bundle defects (D), disrupted cortical actin (E), and defective ADI (F) in *Jabba*-/+; *dCOX1*-/+ compared to single heterozygous controls.

This notion of two distinct pathways is further reinforced when the spatial distribution of LDs is examined. In wild-type and *ATGL* mutant S10B follicles, LDs are evenly distributed throughout the nurse cell cytoplasm (Supplemental Figure S3A-B’). In *Jabba* and *dCOX1* mutants, LDs are unevenly spaced, clustered in some regions of the cytoplasm and depleted from others (Supplemental Figure S3C-E). Similarly, treatment with the COX inhibitor aspirin results in both actin remodeling defects and LD clustering (Supplemental Figure S3F-H”). These LD distributions are reminiscent of LD clustering observed in *Jabba* mutant embryos (Li *et al*., 2012; Kilwein *et al*., 2023a). While the mechanistic basis for this clustering remains to be determined, these observations provide additional evidence of a functional connection between Jabba and dCOX1 that is not shared with the ATGL pathway.

### Jabba acts downstream of dCOX1 to regulate actin remodeling

To uncover how Jabba and dCOX1 act in the same pathway, we first asked if levels or localization of Jabba and dCOX1 are affected in S10B follicles when the other protein is absent. In *Jabba* mutants, dCOX1 levels are unaffected, as measured by western analysis (Supplemental Figure S4A-B); in addition, immunostaining revealed that dCOX1 is still localized to the endoplasmic reticulum, as in wild-type S10B follicles (Supplemental Figure S4C-D’’’’). However, when dCOX1 is absent, Jabba levels and distribution are altered. By western blot analysis, we detected three major Jabba bands in both wild-type and *dCOX1* mutant follicles. They represent three different splice forms of Jabba, B, G, and H (Stephenson *et al*., 2021). Intriguingly, the ratio of G to B isoforms is reduced in the absence of dCOX1 (Supplemental Figure S4E-F). By immunostaining for all Jabba isoforms, we find that Jabba is unevenly distributed throughout the nurse cell cytoplasm in *dCOX1-/-* nurse cells, reminiscent of the clustering of LDs in this genotype (Supplemental Figure S3D-E). Indeed, Jabba still colocalizes with LDs and thus is presumably displaced together with LDs (Supplemental Figure 4G-H’’’’). These changes to Jabba in the *dCOX1* mutant follicles suggest that certain aspects of Jabba are regulated by PG signaling and that Jabba, in some manner, is downstream of dCOX1.

To determine whether Jabba’s impact on actin remodeling is also downstream of dCOX1 and PGs, we increased Jabba levels two-fold in *dCOX1-/-* follicles (as in Fig. 1) and examined whether this affects the actin cytoskeleton in S10B follicles. Our controls (*2x Jabba* on its own and *dCOX1-/-* on its own) recapitulate our findings from Figures 1 and 2 (Figure 5A-B, D-F). Expressing *2x Jabba* in the *dCOX1* mutant resulted in a significant decrease in the frequency of defective bundles (64% vs 85% compared to *dCOX1-/-*) and a striking reduction in severe bundle defects (19% vs 50%; Figure 5C-D). Cortical actin breakdown was also reduced, with 40% (4% severe) of *2x Jabba; dCOX1-/+* follicles exhibiting disrupted cortical actin compared to 88% (65% severe) of *dCOX1*-/- controls (Figure 5E). Similarly, the total actin defects were suppressed in *2x Jabba; dCOX1-/-* follicles as only 11% exhibited a severe ADI compared to 77% in *dCOX1*-/- control (Figure 5F). Comparable findings were observed when *1.5x Jabba* is expressed in the *dCOX1* mutant (Supplemental Figure S5). Because of this dramatic rescue, we conclude that Jabba works downstream of dCOX1 to promote actin remodeling.

**Figure 5.**
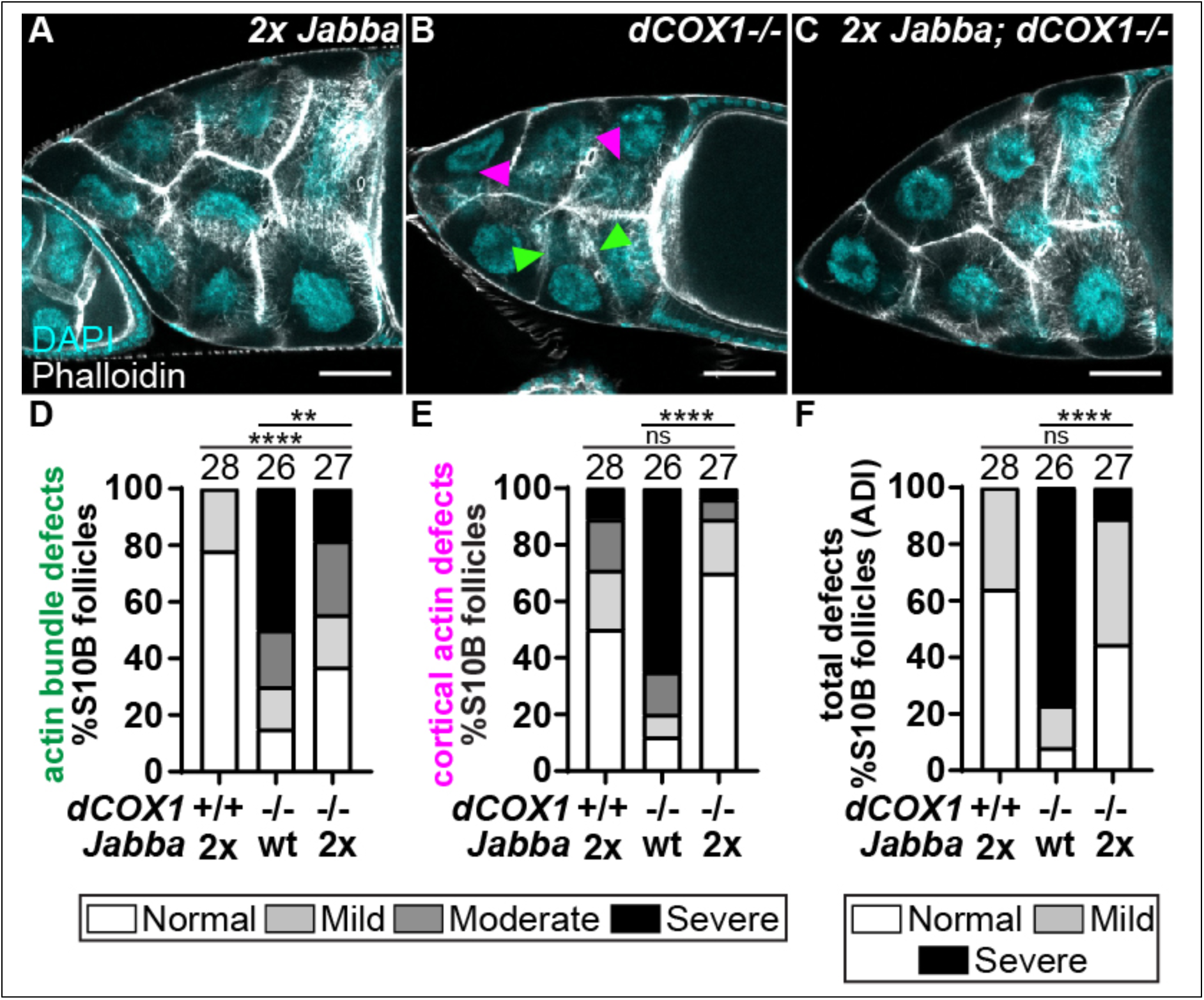
Jabba acts downstream of dCOX1 to promote actin remodeling. (A-C) Maximum projections of three confocal slices of S10B follicles stained for F-actin (phalloidin) in white, and DNA (DAPI) in cyan. Arrowheads indicate instances of actin bundle defects (green) and cortical actin breakdown (magenta). Scale bars = 50µm. (A) *2x Jabba* (two copies of the transgenic genomic *Jabba* [*pJabba*] in a wild-type background). (B) *dCOX1* (*pxt^f01000^/pxt^EY03052^*). (C) *2x Jabba; dCOX1-/-* (*pJabba/pJabba; pxt^f01000^/pxt^EY03052^*). (D-F) Graphs quantifying the frequency of actin defects for the following genotypes: *2x Jabba* (*pJabba*/*pJabba*), *dCOX1*-/- (*pxt^f01000^/pxt^EY03052^)*, and *2x Jabba; dCOX1*-/- (*pJabba/pJabba; pxt^f01000^/pxt^EY03052^*). Actin defects were quantified by scoring the penetrance of actin bundle and cortical actin defects into one of four categories: normal or mild, moderate, or severe defects. Scores were summed and the total binned into one of three total actin defects (ADI) categories: normal, mild defects, or severe defects. For a detailed description of the quantification refer to Materials and Methods. \*\**p<0.01, ****p<0.0001,* Pearson’s chi-squared test. Follicles with increased dosage of *Jabba* form actin bundles and have largely intact cortical actin, though amount of bundles appears increased and the cortical actin appears thicker (A, see Figure 1). Follicles from *dCOX1* mutants, which have wild-type Jabba levels, have disrupted actin bundles and cortical actin breakdown (B, D-F). Overexpression of Jabba in the *dCOX1* mutants suppresses the actin defects, resulting in more normal actin bundle development and cortical actin integrity (C-F).

## DISCUSSION

Our data uncovers a new and unexpected role of the LD-associated protein Jabba in regulating actin remodeling during Drosophila oogenesis. During S10B, overexpression of Jabba drives excessive actin assembly, whereas loss of Jabba results in cortical actin breakdown and severely impaired bundle formation. The actin defects observed when Jabba is lost are strikingly similar to those observed when the LD-associated lipase ATGL or the COX-like enzyme dCOX1 are lost (Tootle and Spradling, 2008; Giedt *et al*., 2023). Dominant genetic interactions and AA toxicity assays reveal that Jabba acts in the same pathway with dCOX1, but not ATGL, to regulate actin remodeling. Genetic studies support that dCOX1, and thereby PG signaling, act upstream of Jabba to drive actin remodeling necessary for follicle morphogenesis and the production of a fertilization competent oocyte (Figure 6).

**Figure 6.**
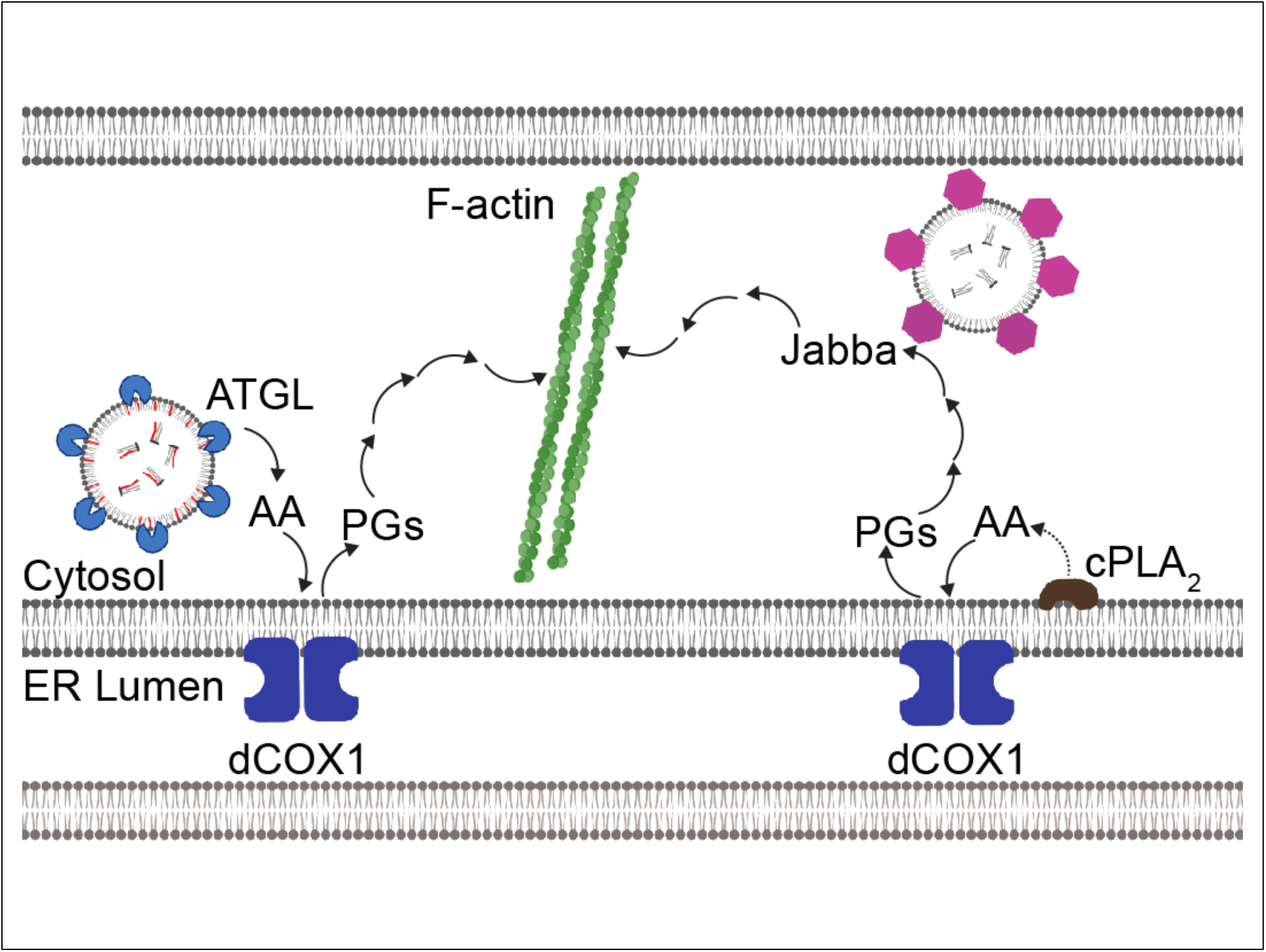
Model of two PG pathways that regulate actin remodeling. Schematic (created in BioRender) depicting two PG-dependent pathways that regulate actin remodeling. On the left, previous studies show that ATGL (light blue) supplies the substrate, AA, for PG production to dCOX1 (dark blue) at the ER. dCOX1 and downstream synthases (not shown) then produce PGs which signal to promote actin (green) remodeling. This paper reveals that the PGs, via an ATGL-independent pathway, regulate the LD-associated protein Jabba (magenta) to promote actin remodeling. In this later pathway, we speculate that cPLA2 (brown) provides the AA for PG production.

Jabba plays multiple roles during *Drosophila* oogenesis. Jabba, via protein-protein interactions, regulates the proteins that associate with LDs (Li *et al*., 2012; Johnson *et al*., 2018). During oogenesis, Jabba binds specific histones to LDs and preserves their use for future embryonic nuclear divisions (Li *et al*., 2012; Li *et al*., 2014; Johnson *et al*., 2018; Stephenson *et al*., 2021). In S14 oocytes, Jabba also prevents LDs from sticking to newly synthesized glycogen granules (Kilwein *et al*., 2023b); if LDs and glycogen are not kept apart, LD allocation to specific embryonic lineages is defective, resulting in delayed development and disruption of redox homeostasis (Kilwein *et al*., 2023b). There are also conflicting reports that Jabba may regulate lipid metabolism during oogenesis: germline specific knockdown of *Jabba* results in mildly decreased neutral lipid accumulation in S10B (McMillan *et al*., 2018), but newly laid embryos from *Jabba* null mutant mothers have triglyceride stores indistinguishable from wild-type embryos (Li *et al*., 2012; Kilwein *et al*., 2023a).

Here we find that Jabba has a new and unexpected function in promoting actin assembly during oocyte development. Overexpression of Jabba results in excess actin assembly, whereas loss of Jabba results in severely impaired actin remodeling. Actin remodeling is energetically intensive; the actin defects observed in the *Jabba* mutants might therefore be due to insufficient energy reserves as a result of reduced neutral lipid stores. However, it is difficult to see how by this mechanism overexpression of Jabba would promote excess actin assembly. Thus, a more likely mechanism whereby Jabba could regulate actin dynamics is by recruiting actin and/or actin binding proteins to LDs. Actin and actin regulators have been found in purified LD preparations in numerous biochemical and proteomic studies (e.g., Fong *et al*., 2001; Pfisterer *et al*., 2017; Bersuker *et al*., 2018; Kilwein and Welte, 2019). LDs might therefore regulate the concentration of these molecules free in the cytosol and thus their availability for actin assembly. If this is the case, our data would suggest that Jabba’s ability to recruit these proteins is regulated by PG signaling. Further, since LDs are highly mobile (Kilwein and Welte, 2019), including in nurse cells (Lu *et al*., 2021), they may deliver actin and its binding partners to sites of actin remodeling within the cell. Intriguingly, the extensive LD clustering in *Jabba* mutants may interfere with their motility and contribute to the actin remodeling defects (Li *et al*., 2012; McMillan *et al*., 2018). While many questions remain, our findings have uncovered a critical role of the LD protein Jabba in regulating actin cytoskeletal remodeling.

Consistent with our findings, there is growing evidence that LDs and their associated proteins regulate actin cytoskeletal remodeling (e.g., Fong *et al*., 2001; Pfisterer *et al*., 2017; Bersuker *et al*., 2018; Kilwein and Welte, 2019). For example, LD-associated proteins bind to the actin regulator actinin to promote the migration and fusion of myoblasts to generate multi-nucleated muscle fibers (Tan *et al*., 2021). In macrophages, many actin regulators are also associated with LDs, including Non-muscle Myosin IIa and Formin-like 1; these proteins regulate actin filament assembly on the LDs, regulating LD dissociation (Pfisterer *et al*., 2017). These studies support that one conserved function of LDs and their associated proteins is to modulate actin dynamics.

In this study and our prior work, we provide the first evidence suggesting that the roles of the LD-associated proteins in actin remodeling are connected to PG signaling. Specifically, we established that during S10B, TAGs within LDs can be used to provide the substrate, AA, for PG production (Giedt *et al*., 2023); substrate release is the rate-limiting step in PG synthesis (Funk, 2001; Tootle, 2013). We found that ATGL cleaves TAGs to release AA, this AA is used by dCOX1 to produce a PG intermediate that is acted on by a downstream synthase to produce PGF_2α_ (Giedt *et al*., 2023). PGF_2α_ then activates a signaling pathway that converges on multiple actin binding proteins to drive actin remodeling necessary for follicle morphogenesis and the production of a fertilization competent oocyte (Groen *et al*., 2012; Spracklen *et al*., 2014; Spracklen *et al*., 2019).

Here we provide evidence that there are two distinct sources of AA needed for PG production during S10B (Figure 6). Through genetic interaction studies we find that Jabba acts independently of ATGL but acts within a distinct PG pathway to promote actin remodeling. Further supporting that Jabba acts independently of ATGL, Jabba has no role in buffering free AA levels unlike ATGL (Giedt *et al*., 2023) and that loss of Jabba or dCOX1, but not ATGL, leads to spatial redistribution of LDs. Finally, overexpression of Jabba suppresses the actin defects in *dCOX1* mutants, indicating Jabba acts downstream of PG signaling. These findings suggest that an ATGL-independent PG synthesis and signaling pathway, in addition to the ATGL-PG pathway (Giedt *et al*., 2023), is required for promoting actin remodeling. We speculate that the ATGL-independent PG pathway that regulates Jabba uses a cytoplasmic phospholipase A2 (cPLA2) to release AA from phospholipids. cPLA2 cleavage of phospholipids is the most well-studied mechanism of providing AA for PG production (Funk, 2001; Tootle, 2013). It is unclear if both pathways of PG production are activated at the same time or sequentially, with one inducing the other in a positive feed forward mechanism, to drive actin remodeling during S10B.

We find that PGs act upstream of Jabba to drive actin remodeling during S10B. The actin defects due to loss of dCOX1 are rescued by overexpressing Jabba. Jabba localization to LDs is unaffected by loss of dCOX1, but the relative levels of the different isoforms of Jabba are altered. Specifically, loss of dCOX1 results in a decrease in Jabba G. This finding suggests that the different isoforms of Jabba may have distinct functions, including during actin remodeling, and may be differentially regulated by PG signaling. Future work is needed to test this possibility.

In addition to altering Jabba isoform balance, PG signaling might regulate Jabba function by other mechanisms. First, it might spatiotemporally control Jabba’s interactions with specific binding partners, including potential actin binding proteins that control actin assembly. Supporting this idea, loss of actin binding protein association with LDs can result in the formation of LD-LD contacts (Pfisterer *et al*., 2017), and both *Jabba* and *dCOX1* mutants exhibit clusters of closely apposed LDs. Second, PG might control post-translational modifications on Jabba which could in turn modulate Jabba’s actin regulatory function or its interaction with actin binding proteins. Indeed, *in vitro*, Jabba can be phosphorylated by Casein Kinase 2 at multiple sites (McMillan *et al*., 2018); additional phosphorylation sites in Jabba depend on the phosphatase Calcineurin (Zhang *et al*., 2019). It will be important to determine if PGs mediate Jabba’s post-translational modifications and whether such modifications are relevant for its role as an actin regulator.

Jabba’s best-described role is to physically anchor histones H2A, H2B, and H2Av to LDs (Li *et al*., 2012; Li *et al*., 2014; Kolkhof *et al*., 2017; Stephenson *et al*., 2021). It is conceivable that PGs manipulate Jabba’s capacity to bind these histones, and changes to free histones levels then alter the gene expression of actin or actin binding proteins. In support of this possibility, *Jabba* mutant embryos accumulate excess H2Av in their nuclei (Li *et al*., 2014). By this mechanism, PGs would utilize Jabba to indirectly alter actin or actin binding proteins, and the altered levels of actin assembly when Jabba is lost or overexpressed may be explained by excess or insufficient histone availability. However, this possibility is unlikely as loss of dCOX1 does not alter the expression of actin or its regulators during oogenesis (Tootle *et al*., 2011; Spracklen *et al*., 2014; Groen *et al*., 2015).

While there is no known mammalian ortholog of Jabba, PG regulation of LD function is conserved across organisms. For example, in white adipose tissue, PGE_2_ signaling regulates LD size to control mitochondrial respiration (Ying *et al*., 2017). In the colon, PG signaling is required for the inflammation-induced increase in LD biogenesis (Heller *et al*., 2016). These studies, in conjunction with our work, lead us to speculate that PG regulation of LDs and their associated proteins is conserved across cell types and organisms.

Together, our results reveal Jabba, a LD-associated protein, is a novel regulator of actin remodeling during Drosophila oogenesis. We find that Jabba’s influence on actin remodeling is not mediated through the upstream ATGL-AA controlled step of PG synthesis, as Jabba and ATGL do not genetically interact, and loss of Jabba does not affect AA toxicity. Jabba and dCOX1 function in a common pathway, and our rescue analysis reveals that Jabba works downstream of dCOX1. Our findings uncover a novel pathway by which a LD protein controls actin remodeling and oocyte development and define a new and unexpected role for Jabba in promoting actin assembly downstream of PG signaling.

## MATERIALS AND METHODS

### Reagents and resources

See Table S1 for information on the reagents used in these studies and Table S2 for the specific genotypes used in each figure panel. All raw data used in this study can be found in Table S3.

### Fly stocks

All stocks used in experiments were maintained on Bloomington standard fly food and at room temperature unless noted. The following stocks were used: *y^1^w^1^* (Bloomington Drosophila Stock Center, BDSC, #1495), *Oregon R* (BDSC, #5), *Jabba^zl01^* (Li *et al*., 2012), *Jabba^DL^*(Li *et al*., 2012)*, bmm^1^* (Gronke *et al*., 2003)*, pxt^EY03052^* (BDSC, #15620), *pxt^f01000^* (Harvard Exelixis Collection, (Thibault *et al*., 2004)), *Jabba RNAi* (TRiP.GL01111, BDSC, #365852), and *oskar GAL4* (generous gift from Anne Ephrussi, European Molecular Biology Laboratory, Heidelberg, Germany (Telley *et al*., 2012); also available at BDSC, #44241). We also generated *pJabba(2), a* genomic transgene of *Jabba*, inserted on the second chromosome. The construct *gJabba* (described in (Johnson *et al*., 2018)) was introduced in the genome via PhiC31 integrase-mediated transgenesis at site 25C6 (Best Gene Inc, Chino Hills, CA); for simplicity, this transgene is referred to as *pJabba* in the text and figures. For RNAi knockdown studies, control and experimental crosses were maintained at room temperature, and progeny were incubated at 29°C for 4-5 days and fed yeast paste daily prior to dissection and staining.

### Immunofluorescence and fluorescent reagent staining

Staining method 1 was used in Figures 1-5, and Supplemental Figures S1, 3F-H’’ and S5. Females were kept with males to promote mating and fed wet yeast paste for 3 days prior to dissection (unless noted otherwise). Ovaries were dissected in room temperature Grace’s medium (Lonza or Corning). Ovaries were fixed in 4% paraformaldehyde in Grace’s medium for 10 minutes at room temperature with rocking. Samples were washed and blocked with antibody wash (1x phosphate buffered saline [PBS], 0.1% Triton X-100, 0.1% bovine serum albumin [BSA]) 6 times for 10 minutes each wash. Follicles were stained with 1 U/mL of Phalloidin AlexaFluor 488 (Invitrogen), Phalloidin Alexa Fluor 568 (Invitrogen) or Phalloidin Alexa Fluor 647 (Invitrogen) in antibody wash overnight at 4°C with rocking. The next day samples were washed 4 times in antibody wash for 10 minutes each, then stained with 1:5000 4′,6-diamidino-2-phenylindole (DAPI, 5 mg/ml) in 1x PBS. Samples were washed in 1x PBS and mounted in 1 mg/mL phenylenediamine in 50% glycerol, pH 9 (Platt and Michael, 1983).

For aspirin treatment studies, ovaries were dissected in fresh IVEM medium (Grace’s medium (Lonza), 2.5% Fetal Bovine Serum (Atlanta Biologicals), 1x Penicillin-Streptomycin (from 100x stock, Gibco)). S10B follicles were isolated and transferred to fresh IVEM medium with 3 mM aspirin or an equivalent amount of ethanol for 1-1.5 hours prior to staining. After the incubation period, the medium was removed, and follicles were fixed in 4% paraformaldehyde in Grace’s medium for 10 minutes at room temperature with rocking. Follicles were stained with 1:5000 Nile red (Sigma) and 1 U/mL Alexa Fluor 647 Phalloidin (Invitrogen) for 2 hours at room temperature. Follicles were washed 2 times in 1x PBS, stained with DAPI (5mg/mL) at 1:5000 in 1x PBS for 10 minutes and washed 2 times in 1x PBS. Follicles were mounted in Aqua-Polymount (PolySciences, Inc.).

Staining method 2 was used in Supplemental Figures S3A-E and S4. Females were housed with males to allow for mating and were fed dry yeast for 2 days at room temperature. Ovaries were dissected in PBS-T (1x PBS, 0.1% Triton X-100) and fixed in 4.1% formaldehyde for 12 minutes at room temperature. Ovaries were washed with PBS-T and forceps were used to isolate S10B follicles. Follicles were blocked overnight at 4°C in ovary block (10% BSA, 0.1% Triton X-100, 0.02% sodium azide in PBS). Follicles were incubated in primary antibodies diluted in ovary block overnight at 4°C. Primary antibodies and concentrations used were rabbit anti-dCOX1/Pxt, 1:1000 (preabsorbed on *dCOX1-/-* ovaries) (Spracklen *et al*., 2014), mouse anti-Calnexin99A, 1:100 (Cnx99A 6-2-1, obtained from the Developmental Studies Hybridoma Bank developed under the auspices of the National Institute of Child Health and Human Development and maintained by the Department of Biology, University of Iowa, Iowa City, IA), and rabbit anti-Jabba, 1:1000 (preabsorbed on *Jabba^DL^* ovaries; (Johnson *et al*., 2018)). Follicles were washed 3 times for 15 minutes each at room temperature with PBS-T. Samples were protected from light and then incubated with secondary fluorescent antibodies diluted 1:1000 in ovary block overnight at 4°C. Secondary antibodies used were goat anti-mouse IgG Alexa Fluor 488 (Invitrogen), and goat anti-rabbit IgG Alexa Fluor 633 (Invitrogen). Samples were washed 3x in PBS-T and then stained with Hoechst 33342 (1 mg/ml, Thermo Fisher Scientific) diluted 1:1000 in ovary block for 20 minutes. For Supplemental Figures 3 and 4, follicles were stained with Nile red (1 mg/ml, Sigma-Aldrich) after immunostaining. Nile red was diluted 1:50 in ovary block for 1 hour at room temperature. Samples were washed 3X in PBS-T prior to mounting in Aqua-Polymount and imaging.

### Fluorescent AA treatment and imaging

Adult female and male flies (to allow for mating) younger than two weeks old were fed dry yeast for 48 hours at room temperature or 24 hours at 25°C in preparation for dissection. Ovaries were dissected in maturation medium (Schneider’s Drosophila medium, Sigma-Aldrich; 15% FBS, Atlanta Biologicals; 10 mg/mL Insulin, Sigma-Aldrich; 1x penicillin/streptomycin, Gibco), and then S10B follicles were isolated and incubated for 15 minutes in maturation medium supplemented with 5 µM fluorescent AA (2-[(7-nitro-2-1,3-benzoxadiazol-4-yl)amino] AA [NBD AA], Avanti Polar Lipids). The medium was removed and the follicles were fixed with 4% paraformaldehyde for 15 minutes. Follicles were stained with LipidSpot 610 (1:100) for 20 minutes, and then washed 3 times for 15 minutes each at room temperature with PBS-T prior to mounting in Aqua-Polymount and imaging.

### Image acquisition and processing

Fixed and stained *Drosophila* follicles were imaged using a Zeiss 700 confocal microscope or Zeiss 880 confocal microscope (Carl Zeiss Microscopy) using a Plan-Apochromat 20x/0.8 working distance (WD)-0.55 M27 objective, a Zeiss 980 (Carl Zeiss Microscopy) using a Plan Apo 20x/0.8 objective, a Leica DM8 Stellaris confocal microscope (Leica Microsystems) using either a HC PL APO CS 20x 0.70 UV objective or a HCX PL APO CS 40x 1.25 oil immersion PH3 UV objective and Leica HyD, or a Leica SP5 confocal microscope (Leica Microsystems) using an HCX PL APO CS 63x/1.40 oil UV objective or HCX PL APO CS 40x/1.25 oil objective and Leica HyD detectors. Images were rotated, cropped, and scale bars were added using FIJI (Abramoff *et al*., 2004), except where noted. As indicated in the figure legends, Adobe Photoshop was used to brighten images. Adobe Illustrator was used to assemble figures.

### Quantification of increased actin assembly and actin defects

Confocal images of phalloidin stained S10B follicles were collected as described. Increased actin was quantified by scanning through genotypically blinded z-stacks of S10B follicles in ImageJ and qualitatively scored to have either normal actin or increased actin bundles and/or thickened cortical actin. Actin bundle and cortical actin defects were scored by scanning through confocal z-stacks of S10B follicles in ImageJ in a genotypically blinded manner. For representative images of follicles and scoring criteria and a detailed description of defect scoring, please refer to Supplemental Figure 1 in Giedt et al. (Giedt *et al*., 2023). Actin bundle defects were scored separately from cortical actin defects. The sum of both scores was binned into one of three categories to determine the ADI. For both quantifications, graphs were created using Prism 10.3.1 (GraphPad Software) and Pearson’s chi-squared analysis with Fisher’s exact test was performed using the rcompanion package (www.rcompanion.org) in R (www.r-project.org).

### Quantification of LD clustering

Confocal images of Stage 10B follicles were scored in a genotypically blinded manner for LD clustering and actin defects. LD clustering was scored based on the penetrance of the phenotype with normal defined as ≤2 nurse cells exhibiting clustering, mild being defined as 3 to 4 nurse cells with clustering, and severe defined as >4 nurse cells with clustering. In some cases, the same follicles were scored for whether they had actin bundle defects and/or disrupted cortical actin as described above. Graphs were created using Prism 10.3.1 and Pearson’s chi squares analysis was performed using R.

### *In vitro* egg maturation (IVEM) assay

For the AA buffering assay (Figure 3), females were maintained on wet yeast paste for 3-4 days prior to dissection. Ovaries were dissected in fresh IVEM medium. Stage 10B follicles were isolated and transferred to clean IVEM medium. For each genotype, 20-30 follicles were distributed between two wells of a 24 well plastic tissue culture dish (Falcon). Either AA (Cayman Chemical) to a final volume of 500µM in 1mL of fresh IVEM medium or an equivalent amount of ethanol as a control was added to each well. Follicles were incubated overnight at room temperature in the dark. The next day, follicles S12 and above were scored as matured. Graphs were created in and statistical analysis was performed using Prism 10.3.1.

### Western blots

Females were housed with males to allow for mating and were fed dry yeast for 2 days at room temperature. Ovaries were dissected in PBS-T (1x PBS, 0.1% Triton X-100) and fixed in 4.1% formaldehyde for 12 minutes at room temperature. Ovaries were washed with PBS-T and forceps were used to isolate S10B follicles. 25-50 S10B follicles were collected per sample and boiled for 20 minutes in 2X Laemmli buffer with 2-mercaptoethanol; both from Bio-Rad. Proteins from fixed samples run at the same molecular weights as fresh samples. Protein samples were run on 10% or 4-15% SDS-PAGE gels (BioRad) and transferred to PVDF membrane for 30 minutes at 80V in Towbin (anti-dCOX1) or 40 minutes at 50V in CAPS (anti-Jabba). Membranes were blocked for 1 hour at room temperature in Odyssey blocking buffer (Licor). Membranes were incubated in primary antibody overnight at 4°C. Primary antibodies were rabbit anti-dCOX1/Pxt (Spracklen *et al*., 2014), 1:5000; mouse anti-α-Tubulin (Millipore Sigma), 1:5000; and rabbit anti-Jabba (Johnson *et al*., 2018), 1:5000. The Jabba antibody was preabsorbed by incubating antibody with membranes containing only *Jabba* null samples (preabsorbed overnight at 4°C). Membranes were washed 2x in Tris buffered saline (TBS)-0.1% Tween 20 for anti-dCOX1/anti-α-Tubulin or 1x PBS-0.1% Tween 20 for anti-Jabba/anti-α-Tubulin. Membranes were protected from light and incubated in secondary antibody for 1 hour at room temperature. Secondary antibodies used were IRDye 800CW goat anti-rabbit IgG (1:10,000, LI-COR) and IRDye 680RD goat anti-mouse IgG (1:10,000, LI-COR). Membranes were washed 2x in TBS/PBS-Tween and then once in TBS/PBS. Membranes were imaged on a LI-COR Odyssey CLx imager and blots were processed and quantified using Image Studio Lite ver 5.2. Graphs were created using Prism 10. Full western blots are shown in Supplemental Figure S6.

## ACKNOWLEDGEMENTS

We thank the Tootle lab and the Welte lab for helpful discussions and careful review of the manuscript and Pakinee Phromsiri with assistance generating *pJabba(2)*. Stocks obtained from the Bloomington Drosophila Stock Center (NIH P40OD018537) were used in this study. FlyBase (release FB2024_04) was used for information on stocks. At the University of Iowa, Information Technology Services – Research Services provided data storage support. This project was supported by the National Institutes of Health (GM116885 and GM144057 to T.L.T., GM102155 to M.A.W.). M.S.G. was partially supported by National Institutes of Health grant T32 CA078586 Free Radical and Radiation Biology, University of Iowa, and I.J.W. was partially supported by National Institutes of Health grant T32 GM144636 Pharmacological Sciences, University of Iowa. J.M.T. was partially supported by National Institutes of Health grants HD114777 and HD10173 to Dr. Mariana Wolfner at Cornell University.

---

**Supplementary Figure S1.**
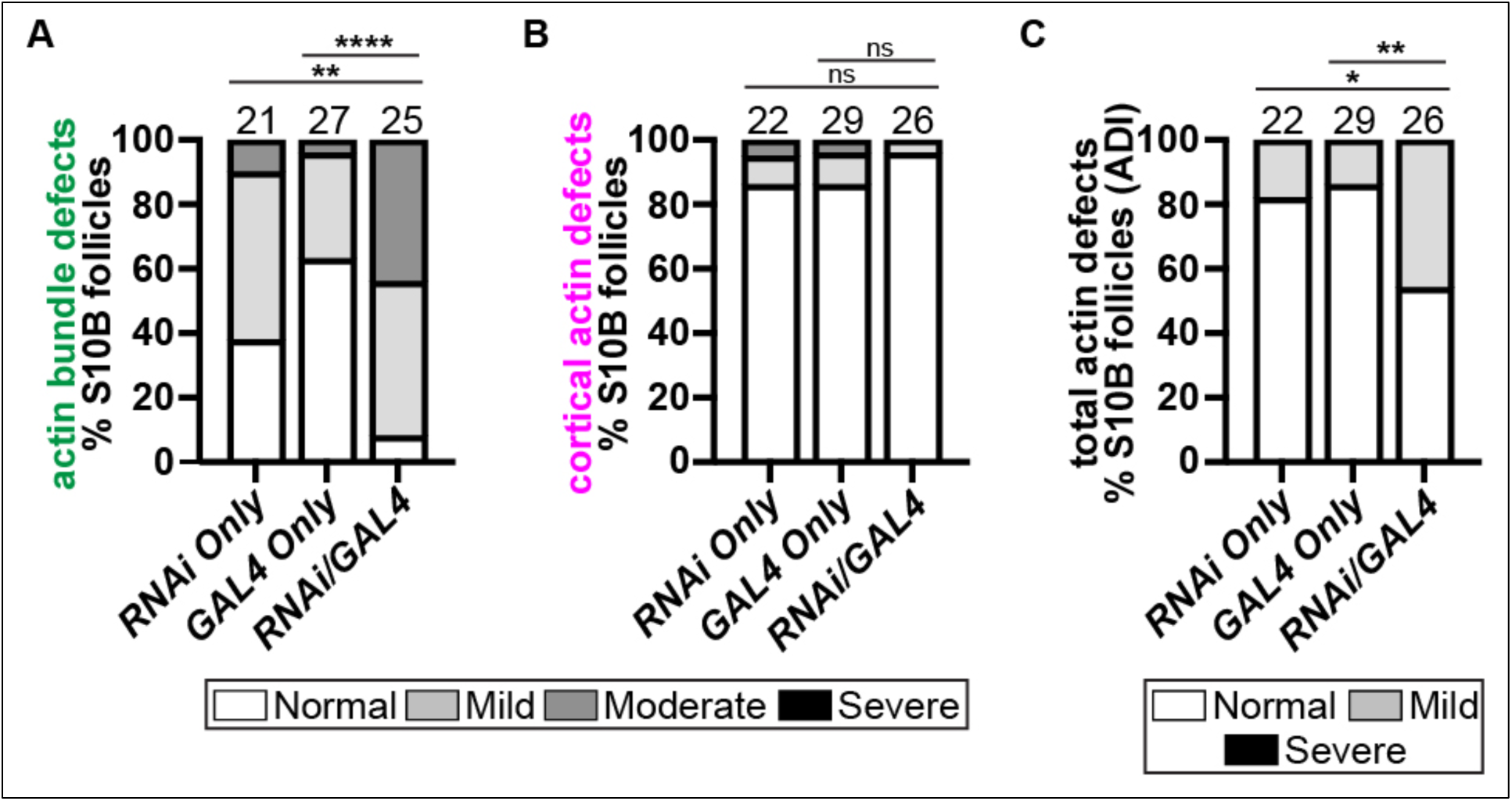
Jabba is required in the germline for normal actin remodeling. (A-C) Graphs quantifying the frequency of actin defects the following genotypes: Jabba RNAi only control (*Jabba RNAi/+*; TRiP.GL01111); GAL4 only control (*oskar GAL4/+*); and germline knockdown of Jabba (*Jabba RNAi/oskar GAL4*). Actin defects were quantified by scoring the penetrance of actin bundle and cortical actin defects into one of four categories: normal or mild, moderate, or severe defects. Scores were summed and the total binned into one of three total actin defects (ADI) categories: normal, mild defects, or severe defects. For a detailed description of the quantification refer to Materials and Methods. ns *p>0.05*, \**p<0.05*, \*\**p<0.01*, \*\*\*\**p<0.0001*, Pearson’s chi-squared test. Germline knockdown of Jabba significantly increases the frequency of moderate actin bundle defects (A), does not impact cortical actin, and decreases the frequency of the normal ADI category compared to controls (C).

**Supplemental Figure S2.**
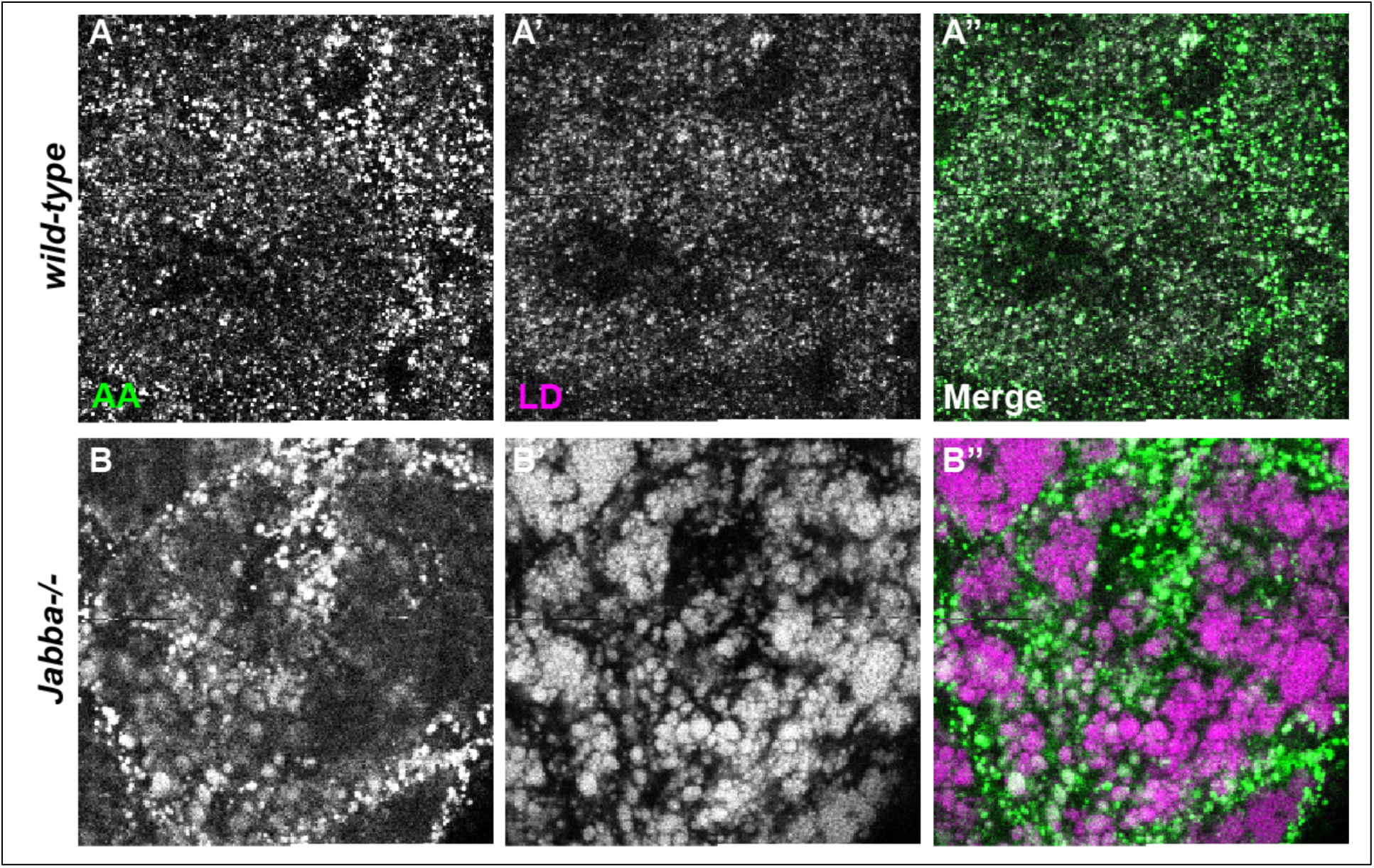
Jabba is not required for AA uptake into LDs. (A-B’’) Single confocal slices of a zoomed in region of the nurse cells in S10B follicles supplemented with arachidonic acid-NBD (AA-NBD; green in merge) and stained for LDs (LipidSpot; magenta in merge). (A-A’’) *wild type* (Oregon R). (B-B’’) *Jabba-/-* (*Jabba^DL^*/*Jabba^DL^*). Scale bars = 10μm. AA is taken up into the LDs of both wild-type and *Jabba* mutant follicles. We note that the highly clustered LDs in the *Jabba* mutant take up less AA.

**Supplemental Figure S3.**
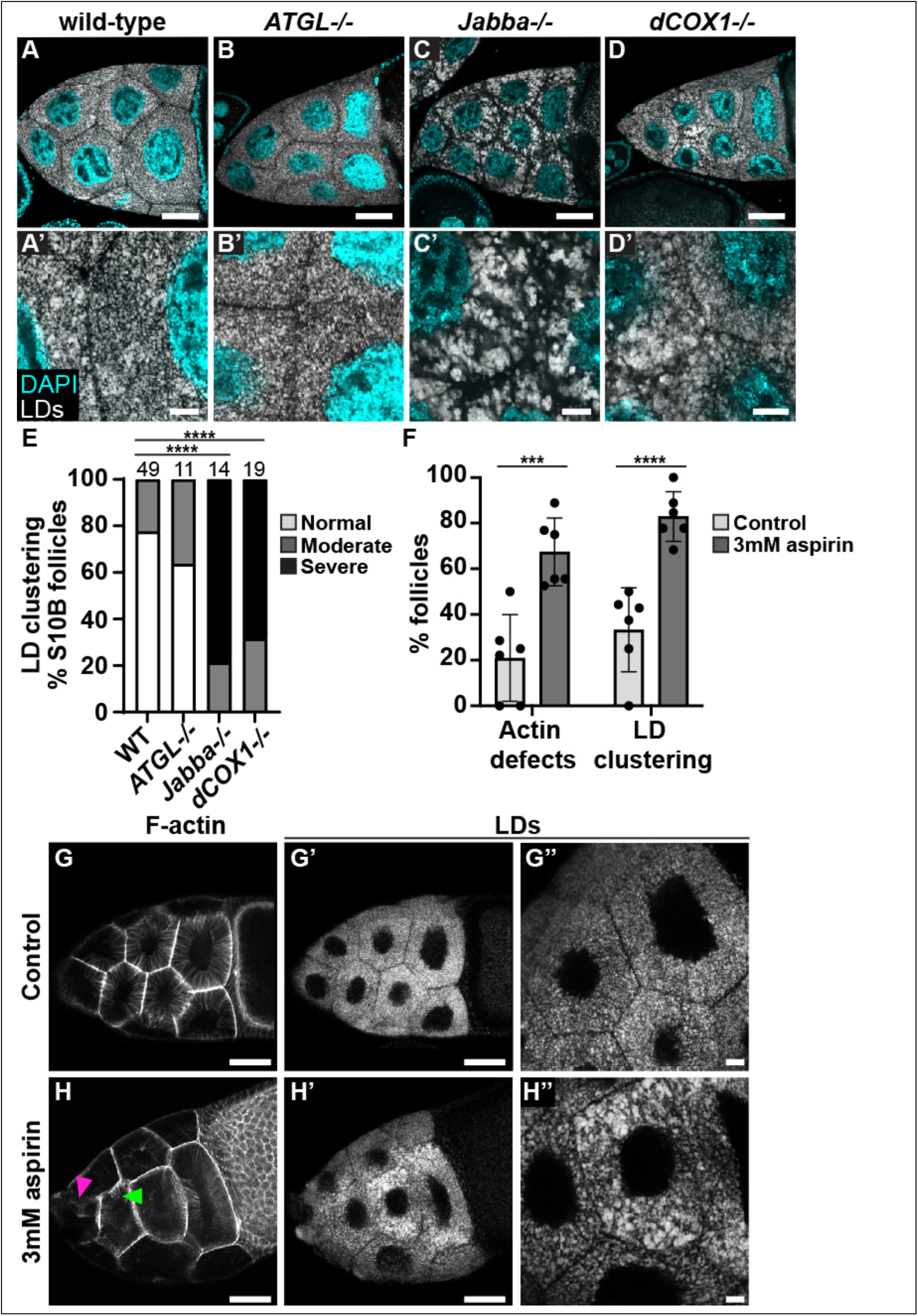
Jabba and PG signaling promote LD dispersal. (A-E’) Single confocal slices of S10B follicles stained for LDs (Nile red) in white and DNA (Hoechst) in cyan; A’-E’ are zoomed in images of A-E. Black boxes were added under the channel labels in A’, and under A’, B’ and D’ to aid in visualization. (A) wild type (Oregon R). (B) *ATGL-/-* (*bmm^1^/bmm^1^*). (C) *Jabba-/-* (*Jabba^DL^/Jabba^DL^*). (D) *dCOX1-/-* (*pxt^f01000^/pxt^f01000^*). (E) Quantification of LD clustering in S10B follicles for the indicated genotypes. Follicles were classified as normal or as exhibiting moderate or severe clustering, and the percentage of each class was calculated. *\*\*\*\*p<0.0001*, Pearson’s chi-squared test. (F) Quantification of actin cytoskeletal and LD clustering defects in control versus 3mM aspirin-treated S10B follicles. Each dot represents the average of a separate experiment; control-treated follicles n=49 and aspirin-treated follicles n=63. Error bars = SD. *\*\*\*p<0.0001, ****p<0.0001*, Sidák’s multiple comparisons test. (G-H’’) Single confocal slices of S10B follicles treated with vehicle (control, EtOH) or 3mM aspirin for 1 hour, and then stained for F-Actin (G, H, phalloidin) or LDs (G’, H’, Nile red) in white; G’’ and H’’ are zoomed in images of G’ and H’. The images in panels G-G’ are on black boxes, and a black box was added under the H’’ label to aid in visualization. Scale bars in A-D, G-G’, H-H’ = 50µm and in A’-D’, G”, I” = 10µm. LDs are evenly distributed in the nurse cells of wild-type and *ATGL* mutant S10B follicles (A-B’, E). Loss of Jabba or dCOX1 results in similar LD clustering/reorganization phenotypes (C-E). Similarly, treatment of S10B follicles with 3mM aspirin, a COX inhibitor, for 1 hour results in both LD clustering (F and H’-H’’ compared to G’-G’’) and actin cytoskeletal defects (F and H compared to G).

**Supplemental Figure S4.**
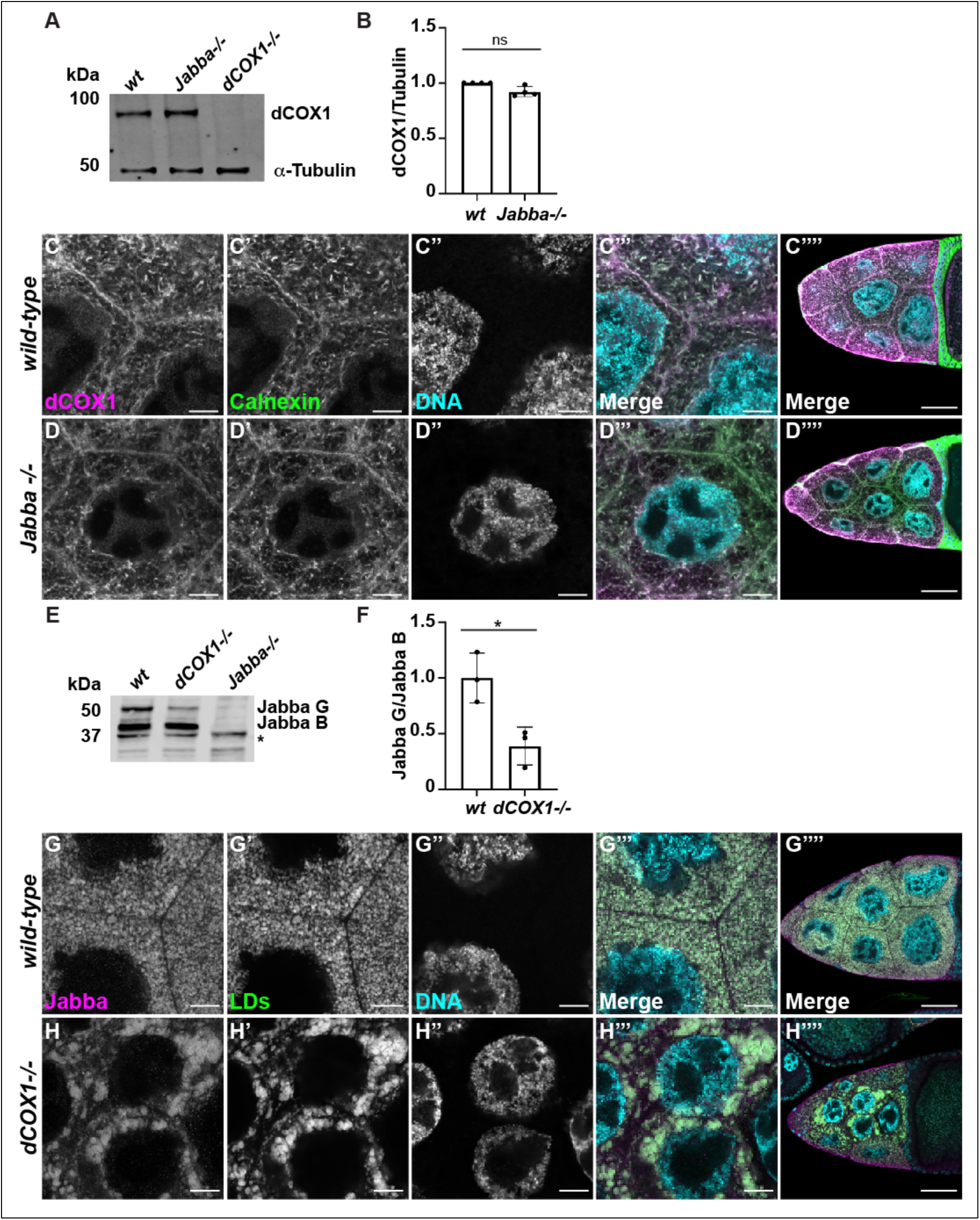
Jabba does not regulate dCOX1 levels or localization, whereas dCOX1 regulates the levels of Jabba isoforms. (A) Western blot of S10B follicles from *wild-type* (*Oregon R)* and *Jabba*-/- (*Jabba^zl01^/Jabba^Zl01^)* ovaries stained for dCOX1 and α-Tubulin (loading control). (B) Quantification of dCOX1 levels in (A). Error bars, SD. p=0.2041. (C-D””) Single confocal slices of nurse cells of *wild-type* (Oregon R; C-C””) and *Jabba-/- (Jabba^zl01^/Jabba^zl01^*, D-D’’’’), stained for dCOX1 (C-D), Calnexin (C’-D’), and DNA (C’’-D’’, Hoechst). Merged image (C’’’-D’’’’): dCOX1, magenta; Calnexin, green; and DNA, cyan. Scale bars in A-D’” = 10µm and C””, D”” = 50μm. Scale bars added in Photoshop. (E) Western blot of wild-type (*Oregon R*), *dCOX1*-/- (*pxt^f01000^*/*pxt^f01000^*), and *Jabba-/-* (*Jabba^zl01^/Jabba^zl01^*) S10B follicles stained for Jabba. Asterisk (*) indicates non-specific band. (F) Graph showing the relative levels of JabbaG/JabbaB in wild-type and *dCOX1*-/- from (E). Error bars, SD. *\*p=<0.0197*, unpaired t-test, two-tailed. (G-H’’’) Single confocal slices of wild-type (*Oregon R,* G-G””) or *dCOX1*-/- (*pxt^f01000^/pxt^f01000^*, H-H””) stained for Jabba (G, H), LDs (G’, H’, Nile red), and DNA (G’’, F”, Hoechst). Merged image (G’’’- H’’’): Jabba, magenta; LDs, green; and DNA, cyan. Scale bars in G-H’” = 10μm and in G””, H”” = 50µm. Scale bars added in Photoshop. Loss of Jabba does not affect dCOX1 expression (A-B) or dCOX1 localization to the ER (C-D””). Loss of dCOX1 reduces the JabbaG/JabbaB ratio compared to wild-type follicles (E-F). Loss of dCOX1-/- does not affect Jabba localization to lipid droplets (G-H””), however the LDs are clustered.

**Supplemental Figure S5.**
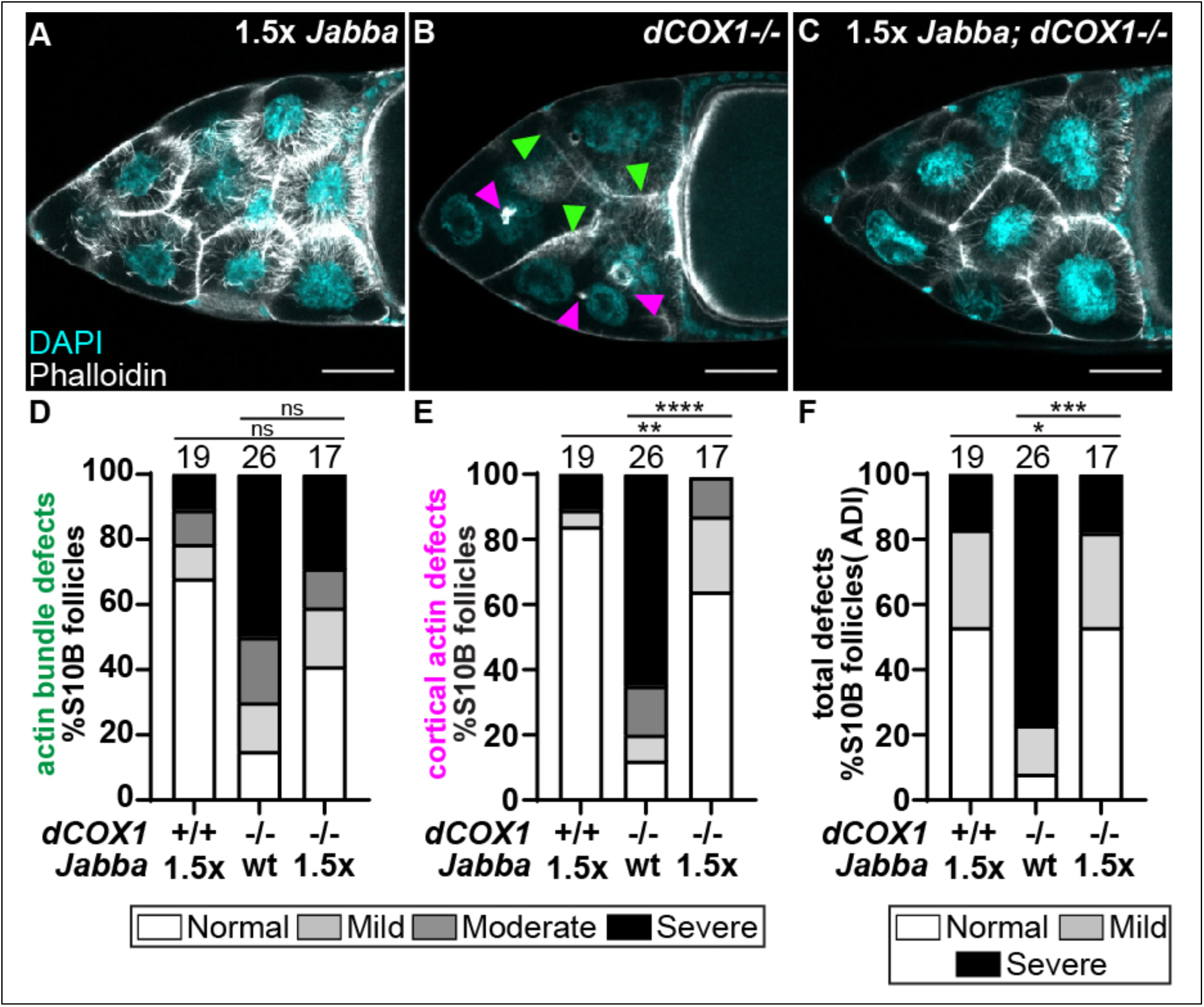
Jabba acts downstream of dCOX1 to promote actin remodeling. (A-C) Maximum projections of three confocal slices of S10B follicles stained for F-actin (phalloidin) in white, and DNA (DAPI) in cyan. Arrowheads indicate instances of actin bundle defects (green) and cortical actin breakdown (magenta). Scale bars = 50µm. (A) *1.5x Jabba* (one copy of the genomic *Jabba* transgene [*pJabba*] in a wild-type background). (B) *dCOX1* (*pxt^f01000^/pxt^EY03052^*). (C) *1.5x Jabba; dCOX1-/-* (*pJabba/+; pxt^f01000^/pxt^EY03052^*). (D-F) Graphs quantifying the frequency of actin defects the following genotypes: *1.5x Jabba* (*pJabba*/*+*), *dCOX1*-/- (*pxt^f01000^/pxt^EY03052^)*, and *1.5x Jabba; dCOX1*-/- ((*pJabba/+; pxt^f01000^/pxt^EY03052^)*. Actin defects were quantified by scoring the penetrance of actin bundle and cortical actin defects into one of four categories: normal or mild, moderate, or severe defects. Scores were summed and the total binned into one of three total actin defects (ADI) categories: normal, mild defects, or severe defects. For a detailed description of the quantification refer to Materials and Methods. Error bars = SD. *\*\*\*\*p<0.0001,* Pearson’s chi-squared test. Follicles with increased dosage of *Jabba* form actin bundles and have largely intact cortical actin, although there appear to be more bundles and cortical actin appears thicker (A, see Figure 1). In contrast, follicles from *dCOX1* mutants, which have wild-type Jabba levels, have disrupted actin bundles and cortical actin breakdown (B, D-F). Overexpression of Jabba in the *dCOX1* mutants suppresses the actin defects, resulting in more normal actin bundle development and cortical actin integrity (C-F).

**Supplemental Figure S6:**
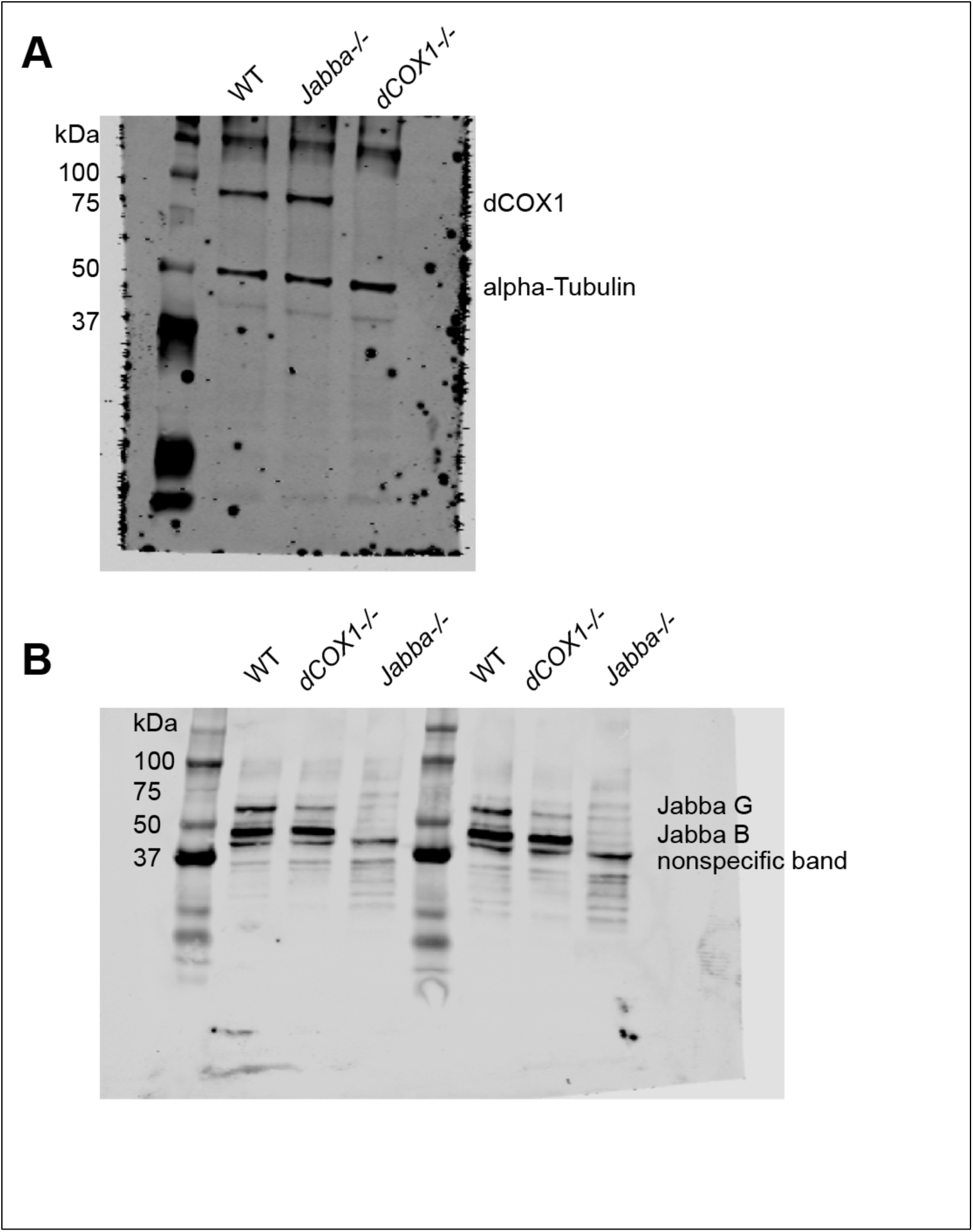
Whole western blots. (A) Whole western blots from Supplemental Figure S3A stained for dCOX1 and α-Tubulin (loading control). (B) Whole western blots from Supplemental Figure S3B stained for JabbaG and JabbaB; nonspecific band indicated. Molecular weight ladders are BioRad Precision Plus Protein Kaleidoscope Standards.

